# Fluctuating selection in a Monkeyflower hybrid zone

**DOI:** 10.1101/2024.06.14.599085

**Authors:** Diana Tataru, Max De Leon, Spencer Dutton, Fidel Machado Perez, Alexander Rendahl, Kathleen G. Ferris

## Abstract

While hybridization was viewed as a hindrance to adaptation and speciation by early evolutionary biologists, recent studies have demonstrated the importance of hybridization in facilitating evolutionary processes. However, it is still not well-known what role spatial and temporal variation in natural selection play in the maintenance of naturally occurring hybrid zones. To identify whether hybridization is adaptive between two closely related monkeyflower species, *Mimulus guttatus* and *Mimulus laciniatus*, we performed repeated reciprocal transplants between natural hybrid and pure species’ populations. We planted parental genotypes along with multiple experimental hybrid generations in a dry (2021) and extremely wet (2023) year in the Sierra Nevada, CA. By taking fine scale environmental measurements, we found that the environment of the hybrid zone is more similar to *M. laciniatus’s* seasonally dry rocky outcrop habitat than *M. guttatus’s* moist meadows. In our transplants hybridization does not appear to be maintained by a consistent fitness advantage of hybrids over parental species in hybrid zones, but rather a lack of strong selection against hybrids. We also found higher fitness of the drought adapted species, *M. laciniatus,* than *M. guttatus* in both species’ habitats, as well as phenotypic selection for *M. laciniatus-*like traits in the hybrid habitat in the dry year of our experiment. These findings suggest that in this system hybridization might function to introduce drought-adapted traits and genes from *M. laciniatus* into *M. guttatus*, specifically in years with limited soil moisture. However, we also find evidence of genetic incompatibilities in second generation hybrids in the wetter year, which may balance a selective advantage of *M. laciniatus* introgression. Therefore, we find that hybridization in this system is both potentially adaptive and costly, and that the interaction of positive and negative selection likely determines patterns of gene flow between these *Mimulus* species.

**Lay Summary:** Early evolutionary biologists understood hybridization, or interbreeding between species, as limiting to adaptation. While recent studies have shown that hybridization is important for adaptation, much remains to be learned about the role of natural selection in maintaining hybridization. We use a repeated transplant experiment in dry and wet years with two closely related monkeyflower species, *Mimulus guttatus* and *Mimulus laciniatus*, and experimental hybrids, to identify how hybridization is maintained. By measuring environmental variables, we found that the hybrid zone is more similar to *M. laciniatus’s* habitat than *M. guttatus’s* in some years. We found that hybrids do equally well as parental species in hybrid zones. Additionally, the drought adapted species, *M. laciniatus,* performed better than *M. guttatus* in both parental habitats, and there was selection for more *M. laciniatus-*like traits in the hybrid habitat. These results suggest that hybridization might introduce drought-adapted traits and genes from *M. laciniatus* in a dry year. In a wet year, first generation hybrids performed better than advanced generation hybrids, possibly due to negative genetic interactions. In summary, we find that hybridization is beneficial and costly, and variation in environmental factors likely determines patterns of hybridization.

## Introduction

Hybridization, or gene flow between species, was seen as a barrier to speciation and adaptation among early taxonomists and speciation researchers (Barton and Hewitt 1985, Arnold 1997). The contribution of gametes to unfit offspring complicated ideas of natural selection introduced by Darwin and muddled the definition of species as groups of reproductively isolated populations (Mayr 1942, Coyne and Orr 2004). This “problem with hybridization” was further corroborated by the discovery of coadapted gene complexes within species, and genetic incompatibilities between species that lead to decreased fitness upon secondary contact. Intrinsic genetic incompatibilities are often masked in first generation hybrids due to heterozygosity, but negative epistatic interactions between alleles are expressed in later hybrid generations (Bateson 1909, Dobzhansky 1936, Muller 1942). Low hybrid fitness in secondary contact could also be due to genotype x environment (GxE) interactions causing extrinsic post-zygotic reproductive isolation (V. Grant 1966, Hatfield and Schluter 1999). While many of these factors do restrict hybridization, evolutionary biologists continue to discover evidence of interspecific gene flow across taxa.

Hybridization was first synthesized as a mechanism of rapid evolution and eventual speciation in the context of adaptation to disturbed habitats (Anderson and Stebbins 1954, Schemske 2000). More recent genomic studies have uncovered hybridization across taxa and habitats as a source of speciation and adaptation through accelerated introgression, increased genetic variation, and heterozygote advantage (Stebbins 1959, Kim and Rieseberg 1999, Coyne and Orr 2004, Abbott, et al. 2013, Mitchell, et al. 2019). One prevailing hypothesis for adaptive introgression is that hybridization is maintained by high fitness in intermediate environments and against hybrids in parental habitats, forming a cline across environmental gradients (Moore 1977). This has been identified in natural systems such as *Artemisia tridentata,* with higher hybrid fitness at an intermediate elevation (Wang, et al. 1997, Miglia, et al. 2005), and in Darwin’s finches, with selection for intermediate beak morphology (Grant and Grant 1996, Grant and Grant 2016). Testing these patterns with experimental hybridization in the wild rather than correlative studies can distinguish between the evolutionary forces that shape patterns of introgression between species (Hatfield and Schluter 1999, Lexer, et al. 2003, Moran, et al. 2021).

Identifying the role of natural selection in shaping population structure is key to understanding the maintenance of hybridization in the wild. Rather than being a consistently directional force pushing populations to an optimum, selection can vary greatly on a spatial and temporal scale (Siepielski, et al. 2013, Wadgymar, et al. 2017, Tataru, et al. 2023). Genetic variation introduced by hybridization can facilitate the persistence of populations in the face of changing conditions, particularly at species’ range edges and during sudden intense environmental changes (Brasier 1995, Grant and Grant 2016). Tension zones are areas where hybridization is maintained by spatial and/or temporal variation in selection, specifically dispersal-selection balance (Key 1968, Bartoń 2009, Gay, et al. 2008, Smith, et al. 2013). When variable environmental conditions interact with novel genetic combinations to produce hybrids with superior fitness to their parental species, this is more aligned with the hybrid novelty model (Arnold 1997). This environment-dependent hybrid advantage, where hybridization was maintained by GxE interactions, is well documented in Louisiana *Iris* species (Johnston, et al. 2001). Identifying the role of environmental variation in shaping hybridization is critical to gaining a deeper understanding of species’ persistence in a changing world.

To test whether hybridization is adaptive in natural hybrid zones we compared two closely related species in the *Mimulus guttatus* (syn. *Erythranthe guttata*) species complex that hybridize in sympatry (Vickery 1964). We conducted reciprocal transplants in hybrid and parental habitats in Yosemite National Park, CA using *Mimulus guttatus, M. laciniatus,* F_1_, F_2_, and reciprocally backcrossed hybrids (Figure 1). We measured phenotypic selection on traits previously identified to be under divergent selection between parental species’ habitats (Ferris, et al. 2014, Tataru, et al. 2023), and fitness differences in two years with drastically different water availabilities. The second year of the experiment (2023) experienced three times the average amount of snowpack, creating extreme conditions and episodic selection. In California’s Mediterranean climate, snowpack and the subsequent amount of snowmelt is critical to water availability in both species’ habitats. In the rocky outcrop habitat of *M. laciniatus* snow melt is ephemeral and dries up quickly, while the deeper soils of *M. guttatus’s* meadows dry out gradually throughout the growing season (Ferris, et al. 2014; Ferris and Willis, 2018; Tataru, et al. 2023). Natural hybrid zones seem to span intermediate habitats, with both meadow and rocky outcrop microhabitats present (Tataru, personal observation).

**Figure 1.**
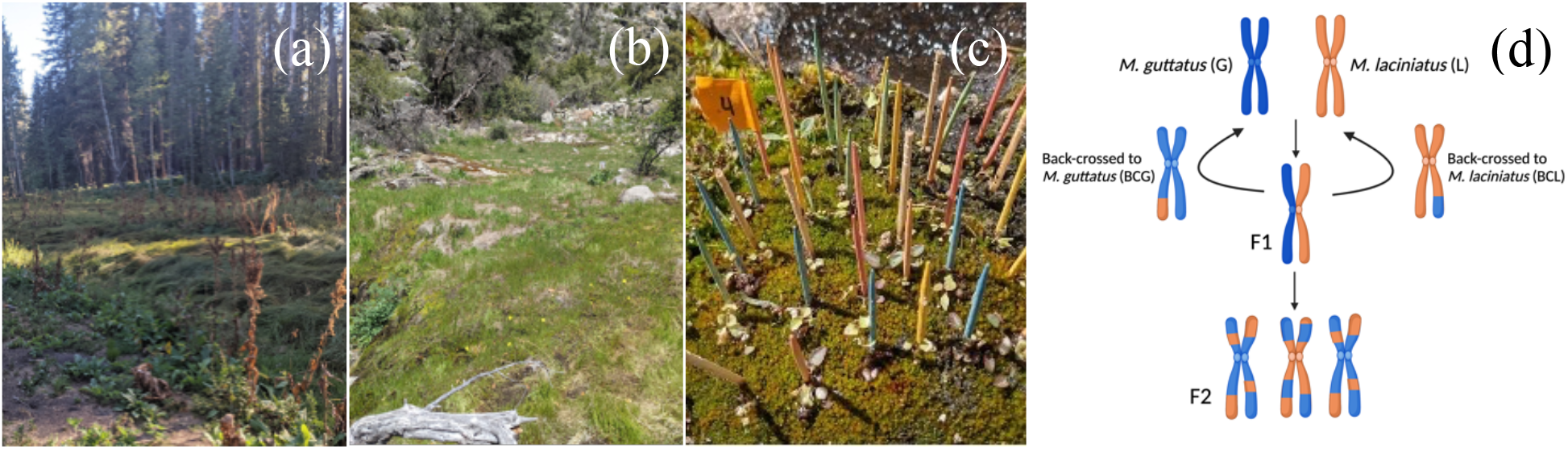
Pictures of (a) M. guttatus parental habitat at the end of the season (b) hybrid zone habitat (c) experimental plot set-up in M. laciniatus parental habitat, and (d) crossing design of six genotypes, with colors indicating parental genetic material (created with Biorender.com).

In our reciprocal transplant experiments we replicated various stages of hybridization likely to be present in a natural hybrid population. Asymmetric introgression due to more frequent backcrossing into one parental species has been well-documented across self-fertilizing vs. outcrossing species pairs similar to *M. laciniatus* (selfer) and *M. guttatus* (outcrosser) (Ruhsam, et al. 2011, Brandvain, et al. 2014, Sianta, et al. 2024). We can test for evidence of adaptive asymmetric introgression by comparing the relative fitness of each direction of experimental backcross hybrids. To our knowledge, there are relatively few experimental hybridization studies of this scale that empirically examine the interaction of natural selection and gene flow in naturally hybridizing species (but see Mitchell, et al. 2019; Campbell, et al. 2024). Using repeated reciprocal field transplants and fine-scale environmental measurements, we aim to answer the following questions: (1) Is the hybrid zone more ecologically similar to the habitat of one species or another? (2) Is gene flow between species advantageous in natural hybrid zones between *Mimulus* species? (3) How does temporally fluctuating selection influence the fitness of hybrids? (4) Which traits are advantageous and under selection in hybrid versus parental habitats?

## Methods

### General Reciprocal Transplant Design

To investigate whether hybridization is adaptive in natural hybrid zones of *M. laciniatus* and *M. guttatus*, we conducted reciprocal transplants in Yosemite National Park, CA, USA over two years with contrasting snowpack levels. We chose transplant sites by the presence of local species or hybrids. The hybrid zone (Figure 1b, Lat: 37.957206, Long: –119.78607, Elevation: 1207 meters) is a moist meadow surrounded by rocky habitat where *M. guttatus*, *M. laciniatus,* and natural hybrids co-occur. The *M. guttatus* site (Figure 1a, Lat: 37.756032, Long: – 119.803024, Elevation: 1841 meters) is a mesic meadow where native *M. guttatus* grows along a seep. Finally, the *M. laciniatus* site (Figure 1c, Lat: 37.85241103, Long: –119.441278, Elevation: 2562 meters) is a granite outcrop with native *M. laciniatus* growing on shallow rocky soils and moss fed by ephemeral snowmelt. The first year of the transplant, 2021, was a drought year with 59% April 1^st^ average snowpack in the Sierra Nevada, CA (CDEC 2024). Experiments ran from April 11^th^ – August 2^nd^, 2021. The second year, 2023, experienced 245% average snowpack (CDEC 2024) and experiments ran May 7^th^ – November 14^th^.

### Environmental Variation Within and Between Habitats

To determine environmental variation among our hybrid and parental sites, we took fine scale environmental measurements at each site across the growing season in both years. We measured soil moisture with a Dynamax SM150 Soil Moisture sensor, surface temperature using a laser thermometer, and light measurements with an Apogee MQ-200X Sunlight Quantum Meter at each block and site every week. We conducted all analyses in R Statistical Software (v4.2.1, R Core Team 2022). To test the effect of environmental variables on survival we used linear mixed effects models in the R package *nlme* (Pinheiro et al. 2023) with plant survival as the dependent variable, and soil moisture, surface temperature, light levels, date, and site as dependent variables, and block as a random effect. We used the R package *MuMin* to conduct model selection (Bartoń 2009). We measured survival as the proportion of plants surviving in each block at the time of each environmental measurement.

To test whether the hybrid zone was either environmentally intermediate or more similar to one of the parental species’ sites, we tested for differences between the slopes of soil moisture decrease over time by running linear mixed effects models in *nlme* (Pinheiro, et al. 2023) with soil moisture as a dependent variable and site, time, and their interaction as independent variables. We then used the R package *emmeans* to compare slopes of linear regressions (Lenth 2016). To quantify differences in the shape of seasonal soil moisture curves, we ran principal components analyses (PCAs) on each year separately. Using the method outlined in Cuthill et al. (1999) we binned measurements by week and input each week as a variable in the PCA. PC1 represents overall variation in total soil moisture values, while PC2 and PC3 represent differences in shape of soil moisture curves over time. We conducted ANOVAs with post-hoc Tukey tests on PC axes (dependent variable) with site and block as independent variables.

### 2021 Reciprocal Transplant

To identify if hybrids have a fitness advantage in hybrid zones we conducted a reciprocal transplant of parental species, F_1_, F_2_, and back-crossed hybrids in hybrid and parental habitats. We reciprocally crossed 25 genotypes of field collected *M. guttatus* (HG) and *M. laciniatus* (HL) from the natural hybrid population to make 50 unique F_1_ crosses. We backcrossed and self-fertilized each F_1_ to create backcross-*M. guttatus* (BCG), backcross-*M. laciniatus* (BCL), and F_2_ hybrids. We used these six genotypic categories (see Figure 1d) in a reciprocal transplant experiment in *M. guttatus*, *M. laciniatus*, and hybrid environments to test whether hybrids have a fitness advantage in the hybrid zone, and decreased fitness in parental environments (Figure 2). We staggered the timing of planting and transplanting so that our experimental seedlings would be at the same developmental stage as native *Mimulus* in each site (hybrid site April 11^th^; *M*. *guttatus* site April 28^th^; *M. laciniatus* site May 26^th^). We stratified soaked flats with seeds at 4C for 14 days, and then germinated plants for one week in growth chambers at UC Davis. We transplanted seedlings at the cotyledon stage into 100 randomized blocks of 36 plants each (six per genotype) at each field site. Plants were approximately one inch apart and native *Mimulus* was removed from blocks. Due to limitations with germination, the total number of individuals at each site varied (Table S1; Hybrid: 3,498 total, *M. guttatus*: 3,372 total, *M. laciniatus*: 2,358 total). To account for transplant shock, we replaced any individuals that died three days after planting.

**Figure 2.**
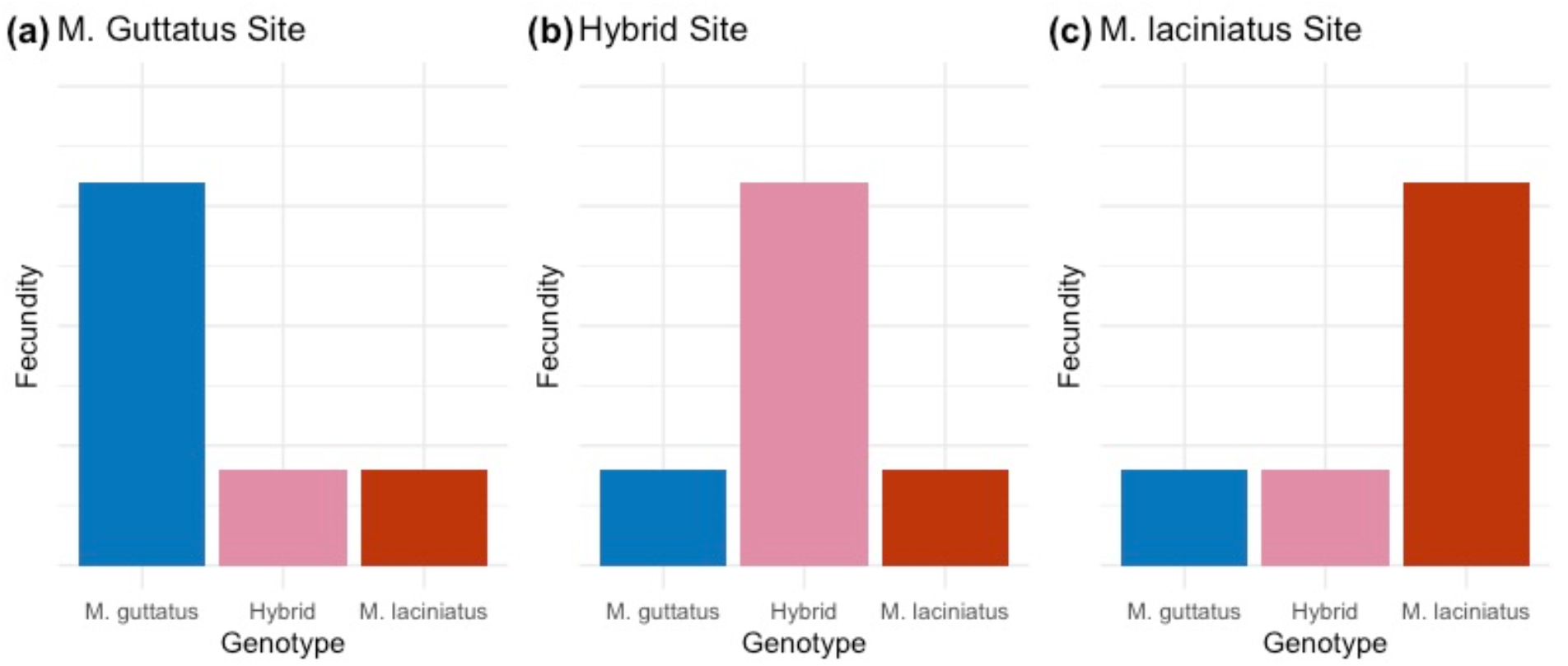
Predicted fecundity in each site given hypothesized fitness advantage of hybrids in hybrid sites, and local adaptation of each species in its native habitat. Plant genotypes are grouped by site, and genotype is indicated by color. Predictions show higher fitness of individuals with *M. guttatus* genes (blue) in the *M. guttatus* habitat (a), higher hybrid (pink) fitness in the hybrid habitat (b), and higher fitness of individuals with *M. laciniatus* genes (red) in the *M. laciniatus* habitat (c).

We monitored sites every three days for timing of first flower, plant height, stigma-anther separation, flower width, leaf measurements, and herbivory. We measured herbivory as presence or absence of damage, and analyzed differences in herbivory between years, sites, and genotypes, and its effect on fitness (survival and fecundity), using a two-way ANOVA. We collected first true leaves on the day of first flower, digitally scanned them, and measured leaf area and lobing index through the program Image J (described in Ferris, et al. 2015). Once plants senesced, we collected all fruits and counted total seed number for each plant which we used as our lifetime fitness metric. We calculated average genotypic fecundity as total seed number divided by the total number of plants planted for each genotype in each habitat.

### 2023 Field Transplant

To examine the role that temporal variation in selection plays in this system, we performed another reciprocal transplant in 2023. We added local genotypes of *M. guttatus* and *M. laciniatus* in addition to the parents of our experimental crosses. In 2023 hybrids were created from one inbred genotype of each parent (*M. laciniatus* maternal x *M. guttatus* paternal). Parental genotypes were outcrossed to control for inbreeding depression (Hall and Willis 2006). We created eight experimental genotypes: Hybrid zone *M. guttatus* (HG), Hybrid zone *M. laciniatus* (HL), local *M. guttatus* (LG), local *M. laciniatus* (LL), F_1_, F_2_, BCL, and BCG. Plants were stratified at UC Merced using the same methodology as the 2021 transplant (see above). We planted 75 blocks of 16 individuals (2 per genotype) at each site (hybrid site May 7^th^ 2023; *M. guttatus* site June 7^th^ 2023; *M. laciniatus* site July 13^th^ 2023). Due to limitations with germination, total number of individuals at each site varied (Table S1; Hybrid: 1,102 total, *M. guttatus*: 990 total, *M. laciniatus*: 1,112 total). Due to logistical constraints, we only measured flowering time phenotype in our 2023 transplant. All other transplant methods were identical to the 2021 transplant (see above).

### Genotypic Selection Statistical Methods

We identified fitness trade-offs between hybrid and parental genotypes at each site across years using linear mixed effects models in *nlme* (Pinheiro, et al. 2023). We tested for genotype by environment interactions using each standardized phenotype as the dependent variable, site (E), genotype (G), and their interaction (GxE) as independent variables, and position nested in block as a random effect. All traits for models were standardized to the mean of 0 and standard deviation of 1 (Kingsolver, et al. 2001). We tested for differences in genotype fitness (survival and fecundity) by calculating mean fitness for every genotype and identified 95% confidence intervals using bootstrap estimates with 1000 repetitions with replacement in the R package *boot* (Davison and Hinkey 1997, Canty and Rippley 2024).

### Phenotypic Selection Statistical Methods

To examine the strength of selection on standardized traits we used zero-truncated poisson and negative binomial linear mixed effect models that account for overdispersion of zeros in the seed set data in the R package *glmmTMB*, with position nested in block as a random effect (Brooks, et al. 2017, Mitchell-Olds and Shaw 1987, Lande and Arnold 1983). We identified best fit models using *MuMin* (Bartoń 2009). In 2023, models only include flowering time. We ran phenotypic selection models in each site on 1) each genotype separately, 2) hybrid genotypes combined, and 3) all genotypes combined. To examine phenotypic correlations among traits in 2021 we ran a correlation analysis using the R package *corrplot* (Wei and Simko 2021).

## Results

### Q1: Hybrid environment fluctuates between being intermediate to more like M. laciniatus’s habitat

Soil moisture was critical for plant survival to flowering in each habitat and year. Soil moisture, time, site, and their interactions all had significant effects on plant survival across years (Figure 3; *p* < 0.0001). Soil surface temperature and ambient light levels were not in best fit models for plant survival in either year (Table 1). Combining environmental data across years, we found a significant effect of year, and the interaction of soil moisture, year, and site on plant survival (Table 1, *p* < 0.0001). This indicates that soil moisture had a strong effect on plant survival and varied over space and time in both a season and over years. There was higher proportion survival to flowering for every genotype in the wetter year (Figure S1).

**Figure 3.**
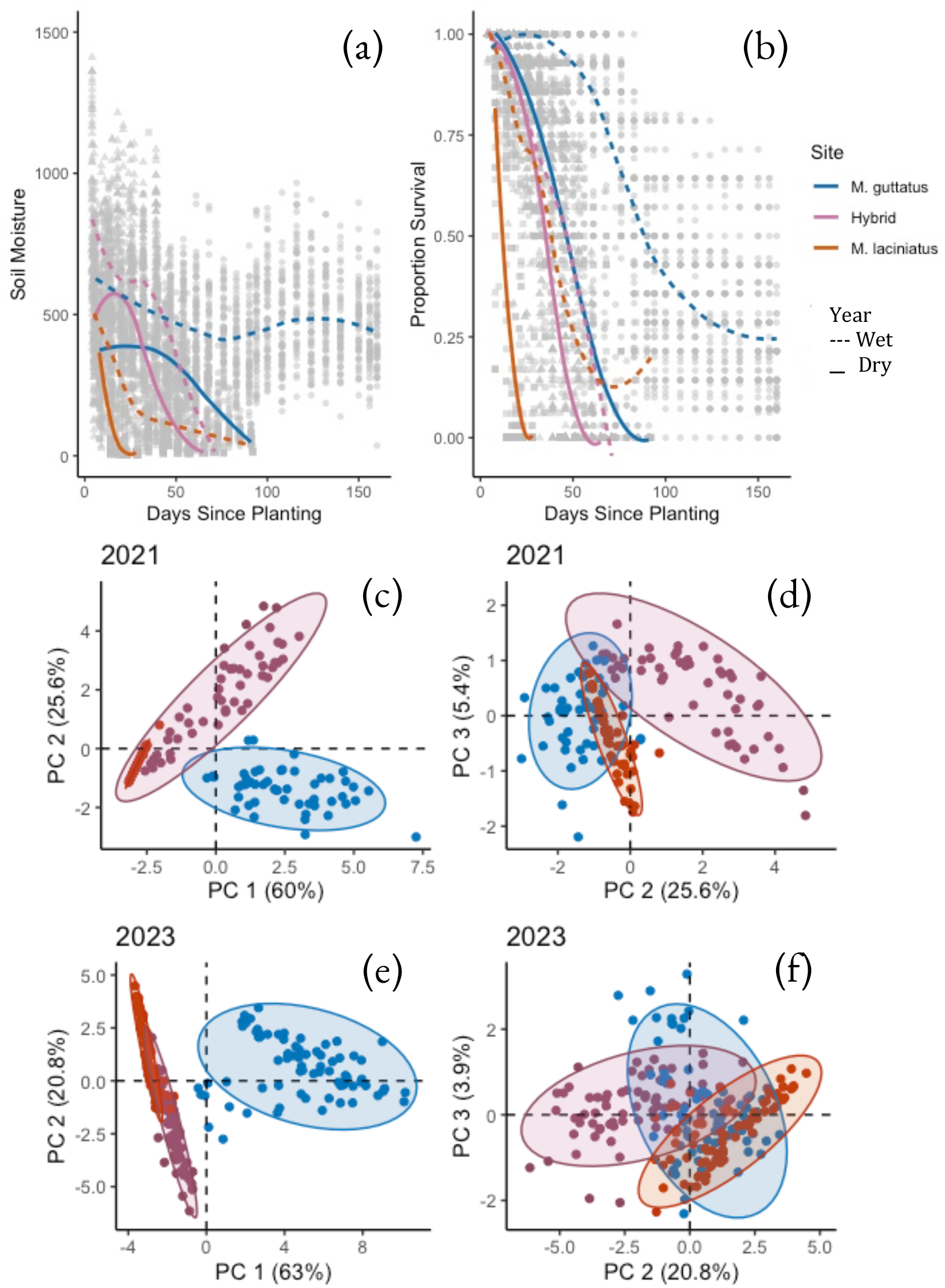
Soil moisture (a) and plant survival (b) decrease over time in *M. guttatus* (blue), hybrid (pink), and *M. laciniatus* (orange) sites. Solid lines connect weekly site means in 2021 and dotted lines connected weekly site means in 2023. Principal component analyses of soil moisture decrease over time in the dry year (c, d) and wet year (e, f). PC1 indicates differences in total soil moisture between site (c, e), while PC2 and PC3 indicate differences in shape of soil moisture curves between sites (d, f).

**Table 1.**
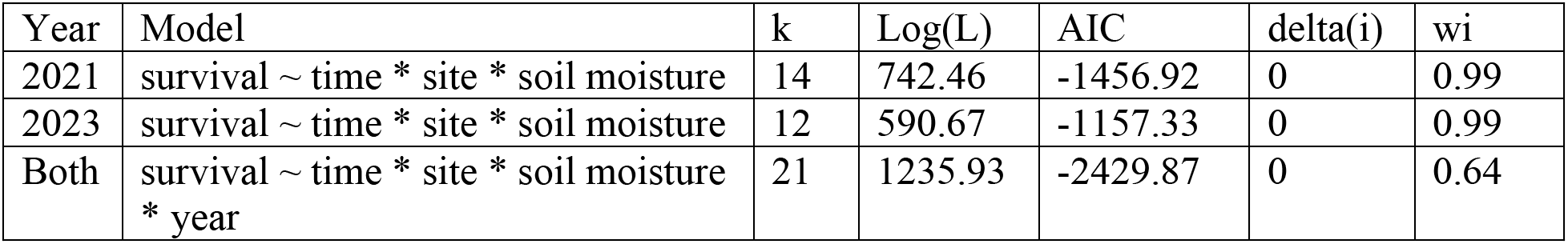
Best fit linear mixed effect models for plant survival with all genotypes combined.

| Year | Model | k | Log(L) | AIC | delta(i) | wi |
| --- | --- | --- | --- | --- | --- | --- |
| 2021 | survival ~ time * site * soil moisture | 14 | 742.46 | -1456.92 | 0 | 0.99 |
| 2023 | survival ~ time * site * soil moisture | 12 | 590.67 | -1157.33 | 0 | 0.99 |
| Both | survival ~ time * site * soil moisture * year | 21 | 1235.93 | -2429.87 | 0 | 0.64 |

To identify whether the hybrid site was intermediate to the two parental sites, we tested for significant differences in slope and shape of soil moisture decrease over time since planting (Figure 3). In both years the slope of hybrid site soil moisture decline was more similar to the *M*. *laciniatus* site (Pairwise comparisons 2021: *t*(1587) = 5.227, *p* < 0.0001; 2023: *t*(2732) = –5.368, *p* < 0.0001) than the *M. guttatus* site (Pairwise comparisons 2021: *t*(1587) = 15.731, p < 0.0001; 2023: *t*(2732) = 31.894, *p* < 0.0001). ANOVAs of soil moisture curve PCAs showed significant differences between all sites in each year, with variation in total soil moisture (PC1) in the hybrid zone intermediate to parental sites in the dry year (Table S2; Figure 3c), and more similar to the *M. laciniatus* site in the wet year (Table S2; Figure 3e). Soil moisture curve shape (PC2) was more similar in parental sites than the hybrid site in both years (Table S2; Figure 3d&f). Therefore, the hybrid zone seems more similar to *M. laciniatus’s* habitat in overall soil moisture levels, but unique from parent species in patterns of seasonal soil moisture decrease.

Herbivory had a significant effect on survival (dry year *f* = 41.72, *p* < 0.0001; wet year *f* = 14.2 *p* = 0.000167) but not fecundity in both years. Proportion herbivory varied significantly by site, year, genotype, and the interaction of all factors (Table S3). The wet year had higher herbivory than the dry year in all habitats, while the hybrid and *M. guttatus* habitats had higher levels of herbivory than *M. laciniatus* in both years (Table 2). Unlike soil moisture, the hybrid zone seems more like the *M. guttatus* habitat in herbivory levels.

**Table 2.** Proportion of herbivory across all genotypes.

| Site | Year | N | Proportion herbivory | sd | se | ci |
| --- | --- | --- | --- | --- | --- | --- |
| *M. guttatus* | 2021 | 3404 | 0.148 | 0.356 | 0.006 | 0.012 |
| *M. guttatus* | 2023 | 974 | 0.173 | 0.378 | 0.012 | 0.024 |
| Hybrid | 2021 | 3511 | 0.169 | 0.375 | 0.006 | 0.012 |
| Hybrid | 2023 | 1030 | 0.285 | 0.452 | 0.014 | 0.028 |
| *M. laciniatus* | 2021 | 2463 | 0.001 | 0.029 | 0.001 | 0.001 |
| *M. laciniatus* | 2023 | 1112 | 0.043 | 0.203 | 0.006 | 0.012 |

### Q2: Relaxed selection against hybrids in hybrid habitat

There were genotypic fitness differences between habitats in both transplant years. In both years we found no significant difference in fecundity between hybrid and parental genotypes in the hybrid habitat, with the exception of higher F_1_ hybrid having fecundity in the wet year (Figure 4b&e; Tables S1). In both years non-local *M. laciniatus* (HL) and BCL had higher fecundity (Figure 4a&d) and higher survival (Figure S1) than most genotypes in *M. guttatus’* habitat. *M. guttatus* genotypes had the lowest total fecundity across all habitats, with local *M. guttatus* (LG) having even lower fecundity than hybrid zone *M. guttatus* (HG) in 2023 (Figure 4d). However, in that wet year ∼40% of LG individuals survived to the end of the growing season without flowering in the *M. guttatus* habitat (Figure S2). These results indicate relaxed selection against hybridization in the hybrid zone, and possible local maladaptation in the *M. guttatus* habitat.

**Figure 4.**
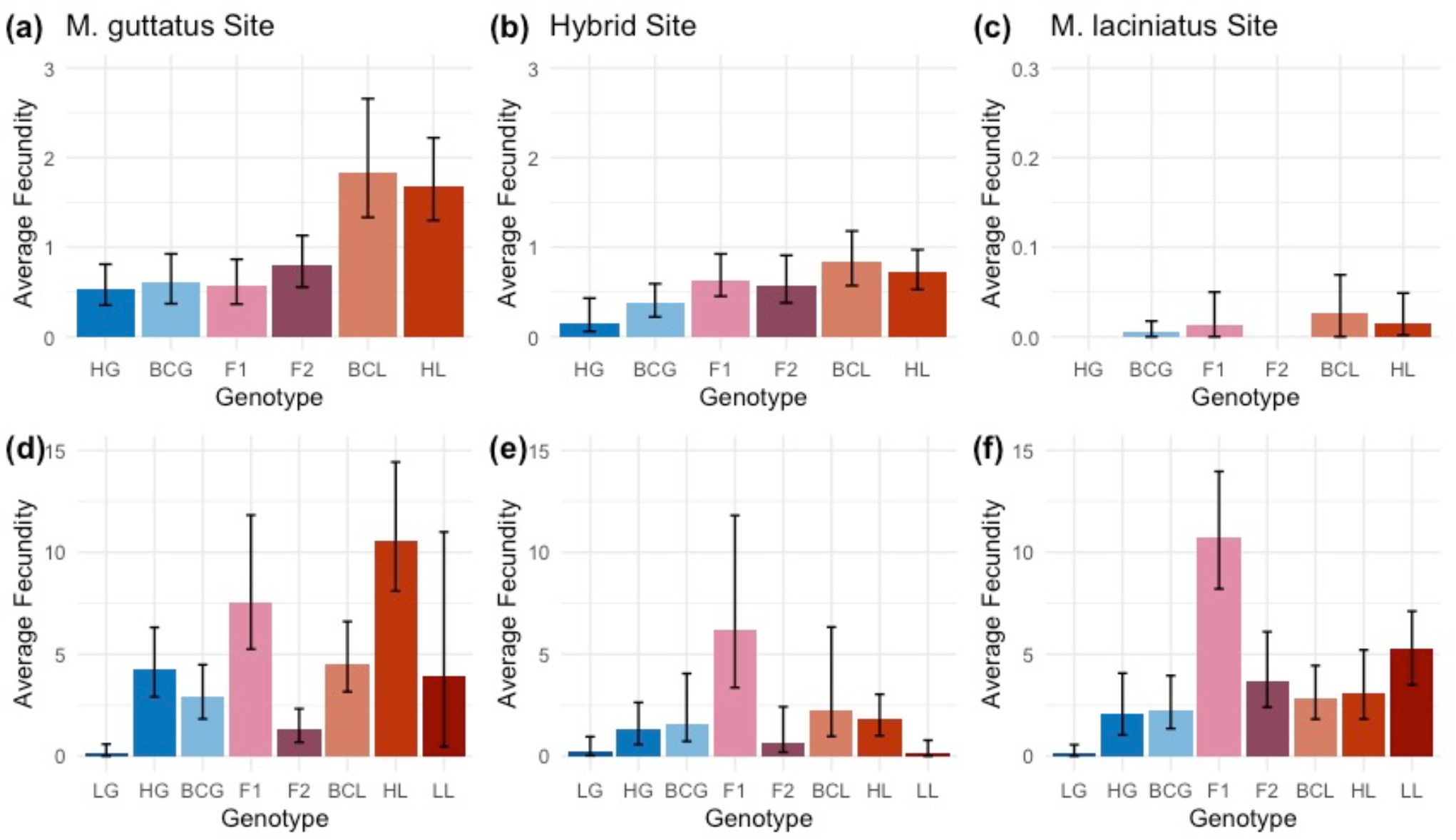
Average lifetime fecundity (total seed number/number of plants planted) per genotype in each site, with sites plotted separately. Upper row (a, b, c) is 2021 sites and lower row (d, e, f) is 2023 sites. Genotypes are, from left to right, local *M. guttatus* (LG), hybrid habitat *M. guttatus* (HG), back-crossed *M. guttatus* (BCG), first generation hybrid (F_1_), second generation hybrid (F_2_), backcrossed *M. laciniatus* (BCL), hybrid habitat *M. laciniatus* (HL) and local *M. laciniatus* (LL). Error bars represent 95% confidence intervals from bootstrap analysis. Values for average fecundity and 95% CI can be found in Supplementary Table 1.

### Q3: Genotypic selection varies temporally

We found temporal variation in selection between our dry and wet transplant years. There was higher fitness across genotypes and sites in the wet year compared to the dry (Figure 4; Figure S1). Patterns of which genotypes performed better also vary between years. In the dry year, genotypes with more *M. laciniatus* genetic background (HL, BCL) had higher fecundity and survival across sites, and most strongly in the *M. guttatus* site (Figure 4; Figure S1).

During the wet year, we saw evidence of a hybrid advantage which was not present in the drier year (Figure 4). F_1_ hybrids had the highest fecundity in both the hybrid zone and *M. laciniatus* habitats, and second highest average fecundity in *M. guttatus’s* habitat (Figure 4d, e, f). F_2_ hybrids had significantly lower fitness than F_1_ hybrids in each habitat, where-as in the dry year they had similar levels of fitness in hybrid and *M. guttatus* habitats (Figure 4). The significant decrease in relative F_2_ fitness in the wetter year is likely due to expression of Bateson-Dobzhansky-Mueller incompatibilities, or BDMIs (Bateson 1909, Dobzhansky 1936, Muller 1942).

### Q4: Phenotypic Selection for M. laciniatus-like traits in the Hybrid Zone

Genotype and environment effected quantitative trait expression in both transplant years. In the dry year we found significant effects of site (E), plant genotype (G), and the interaction of site and genotype (GxE) on flowering time, plant height, stigma-anther separation, and leaf size (Table S4). Site and genotype, but not their interaction, had significant effects on flower size and leaf lobing. In the wet year, site, genotype, and their interaction had a significant effect on flowering time (Table S4).

We conducted phenotypic selection analysis on plants in the *M. guttatus* and hybrid sites in 2021 and on flowering time across sites in 2023. Too few plants survived to flowering to perform selection analyses in the *M. laciniatus* site in 2021. Correlations among traits were relatively weak (r < 0.2, Figure S3) except for size related phenotypes (*ρ* ∼ 0.5) indicating that most traits in our phenotypic selection analysis are weakly correlated. In our combined genotype zero-truncated poisson model we found stronger selection in the hybrid zone than the *M. guttatus* site on most quantitative traits (Figure 5; Table 3). In the hybrid site we found strong selection for earlier flowering, smaller leaves, taller plants, and smaller flowers. These trait values are in the direction of *M. laciniatus* phenotypes, apart from taller plants which is more *M. guttatus*-like. In the *M. guttatus* site we found weak selection for earlier flowering and larger plants (Figure 4). In the hybrid site, the best fit zero-truncated poisson model involved interactions between flowering time, and plant height, leaf size, and stigma-anther separation, as well as an interaction between plant height and stigma-anther separation (Table 3). In contrast, there was only one interaction between flower width and plant height in the *M. guttatus* site (Table 3). These interactions indicate possible correlational selection (Svensson 2023), suggesting stronger correlational selection in the hybrid habitat.

**Figure 5.**
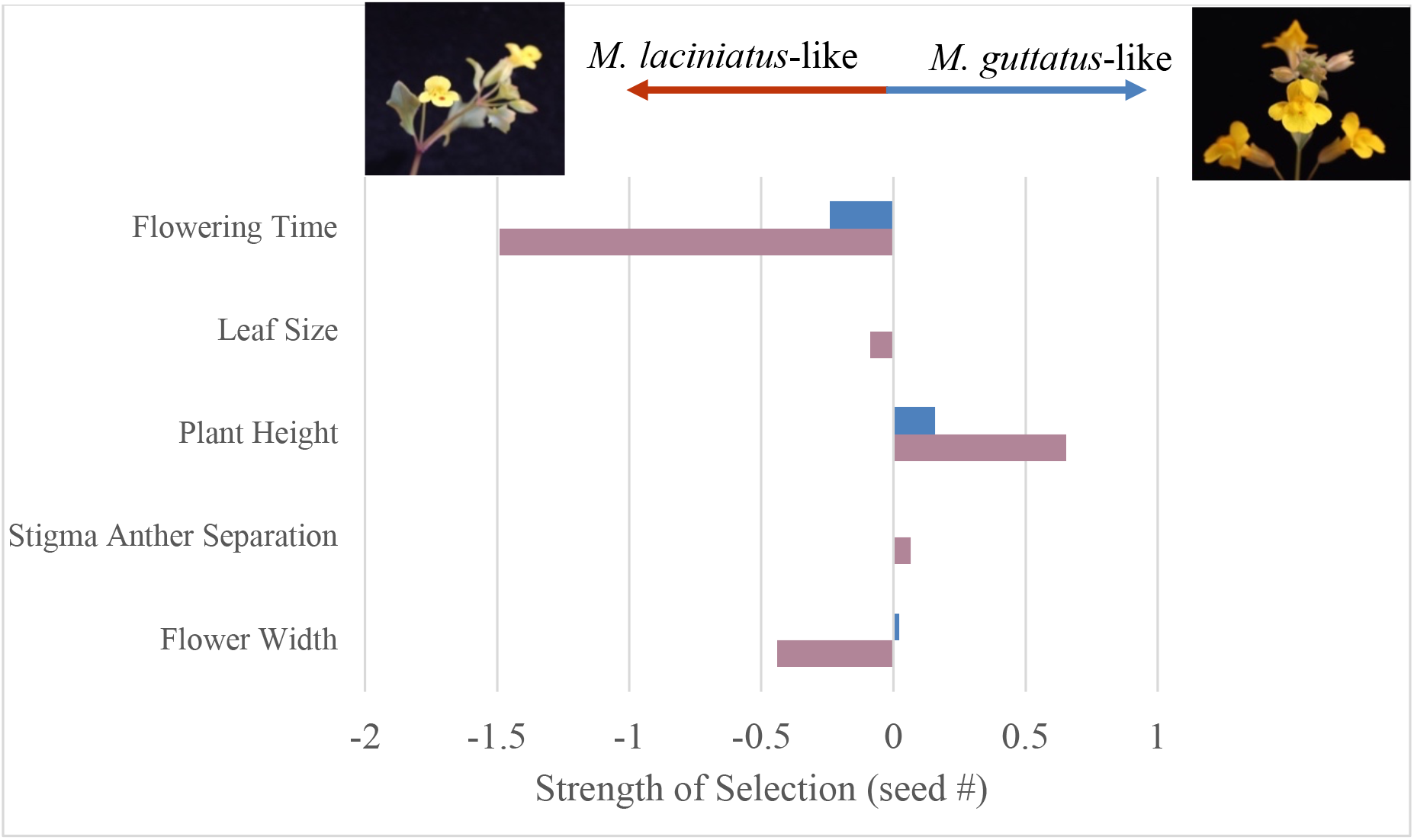
Phenotypic selection analysis in 2021 of all genotypes combined in the *M. guttatus* (blue) and Hybrid (pink) sites. There was no significant selection for leaf roundedness for genotypes combined in either species habitat in 2021. Pictures and arrows above the graph indicate which species’ traits the direction of selection is moving towards, *M. laciniatus* (left) and *M. guttatus* (right).

**Table 3.**
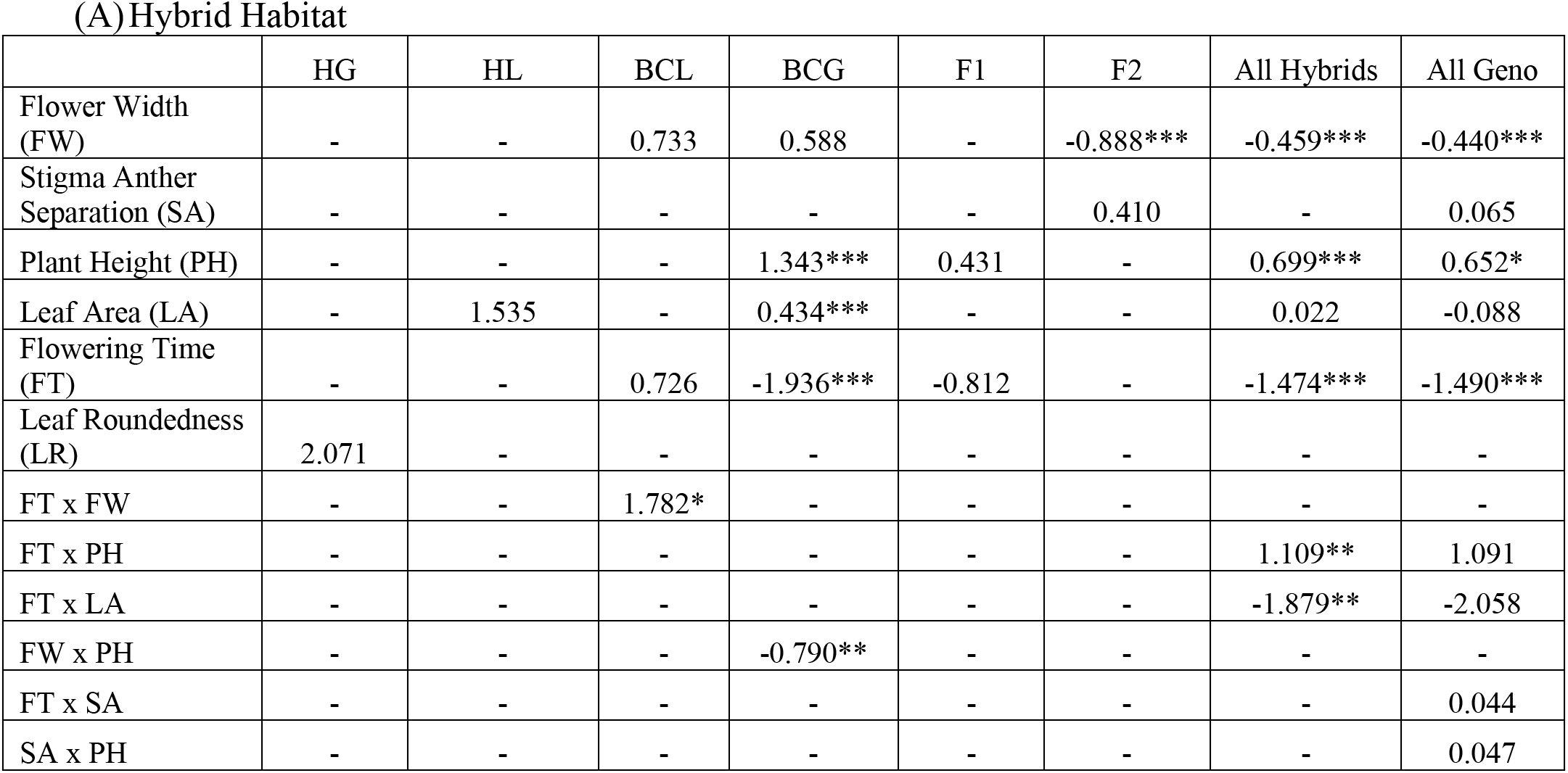
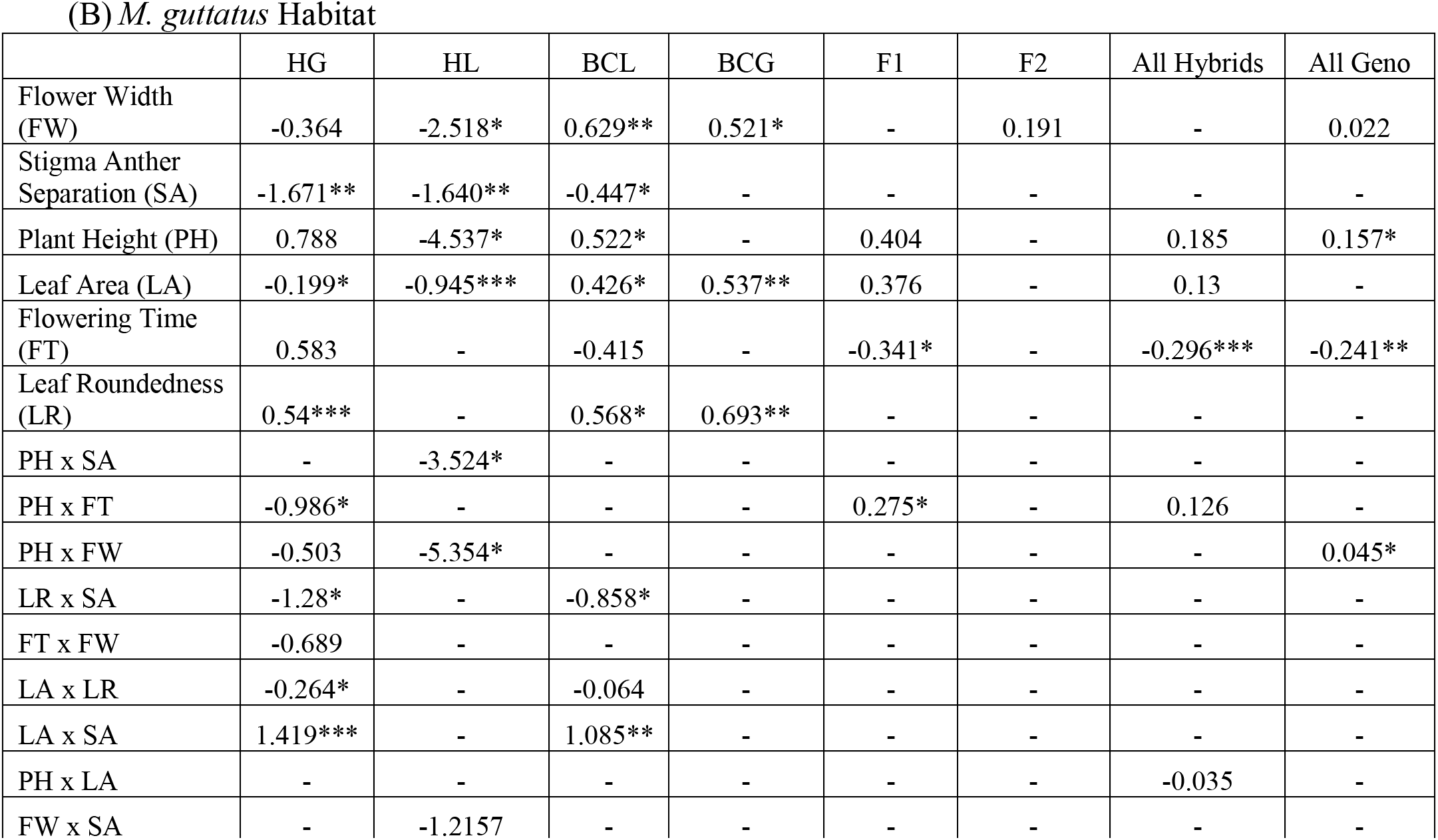
Selection gradients (β) from the best fit zero-truncated poisson models based on 2021 seed number in the hybrid habitat. (A) and *M. guttatus* habitat (B). β indicates strength and direction of selection while asterisks indicate significance: p<.05 *, P<.01**, P< .001 ***.

(A) Hybrid Habitat
|  | HG | HL | BCL | BCG | F1 | F2 | All Hybrids | All Geno |
| --- | --- | --- | --- | --- | --- | --- | --- | --- |
| Flower Width (FW) | - | - | 0.733 | 0.588 | - | -0.888*** | -0.459*** | -0.440*** |
| Stigma Anther Separation (SA) | - | - | - | - | - | 0.410 | - | 0.065 |
| Plant Height (PH) | - | - | - | 1.343*** | 0.431 | - | 0.699*** | 0.652* |
| Leaf Area (LA) | - | 1.535 | - | 0.434*** | - | - | 0.022 | -0.088 |
| Flowering Time (FT) | - | - | 0.726 | -1.936*** | -0.812 | - | -1.474*** | -1.490*** |
| Leaf Roundedness (LR) | 2.071 | - | - | - | - | - | - | - |
| FT x FW | - | - | 1.782* | - | - | - | - | - |
| FT x PH | - | - | - | - | - | - | 1.109** | 1.091 |
| FT x LA | - | - | - | - | - | - | -1.879** | -2.058 |
| FW x PH | - | - | - | -0.790** | - | - | - | - |
| FT x SA | - | - | - | - | - | - | - | 0.044 |
| SA x PH | - | - | - | - | - | - | - | 0.047 |

Table 3. Selection gradients ( $\beta$ ) from the best fit zero-truncated poisson models based on 2021 seed number in the hybrid habitat (A) and *M.
|  | HG | HL | BCL | BCG | F1 | F2 | All Hybrids | All Geno |
| --- | --- | --- | --- | --- | --- | --- | --- | --- |
| Flower Width (FW) | -0.364 | -2.518* | 0.629** | 0.521* | - | 0.191 | - | 0.022 |
| Stigma Anther Separation (SA) | -1.671** | -1.640** | -0.447* | - | - | - | - | - |
| Plant Height (PH) | 0.788 | -4.537* | 0.522* | - | 0.404 | - | 0.185 | 0.157* |
| Leaf Area (LA) | -0.199* | -0.945*** | 0.426* | 0.537** | 0.376 | - | 0.13 | - |
| Flowering Time (FT) | 0.583 | - | -0.415 | - | -0.341* | - | -0.296*** | -0.241** |
| Leaf Roundedness (LR) | 0.54*** | - | 0.568* | 0.693** | - | - | - | - |
| PH x SA | - | -3.524* | - | - | - | - | - | - |
| PH x FT | -0.986* | - | - | - | 0.275* | - | 0.126 | - |
| PH x FW | -0.503 | -5.354* | - | - | - | - | - | 0.045* |
| LR x SA | -1.28* | - | -0.858* | - | - | - | - | - |
| FT x FW | -0.689 | - | - | - | - | - | - | - |
| LA x LR | -0.264* | - | -0.064 | - | - | - | - | - |
| LA x SA | 1.419*** | - | 1.085** | - | - | - | - | - |
| PH x LA | - | - | - | - | - | - | -0.035 | - |
| FW x SA | - | -1.2157 | - | - | - | - | - | - |

When we break up the phenotypic selection analysis by genotype, we find that in both zero-truncated poisson and negative binomial models the hybrid genotypes (F_1_, F_2_, BCL, BCG) in the *M. guttatus* site largely experience selection towards local *M. guttatus* trait values (Figure S4 & S5; Table 3 & S5). In the negative binomial, most genotypes in the hybrid habitat experienced selection towards *M. laciniatus*-like traits (earlier flowering, smaller leaves, smaller plants, and smaller flowers) except for *M. guttatus* (HG) which experienced selection for more *M. guttatus*-like traits: later flowering, larger flowers, and greater stigma-anther distance (Figure S5; Table S5). In the wet year, we found selection for earlier flowering in all sites and models, with the strongest selection for early flowering in the *M. laciniatus* site and hybrid sites (Figure 6h, Table 4). Therefore, in both years we see selection in the hybrid zone for more *M. laciniatus*-like traits.

**Figure 6.**
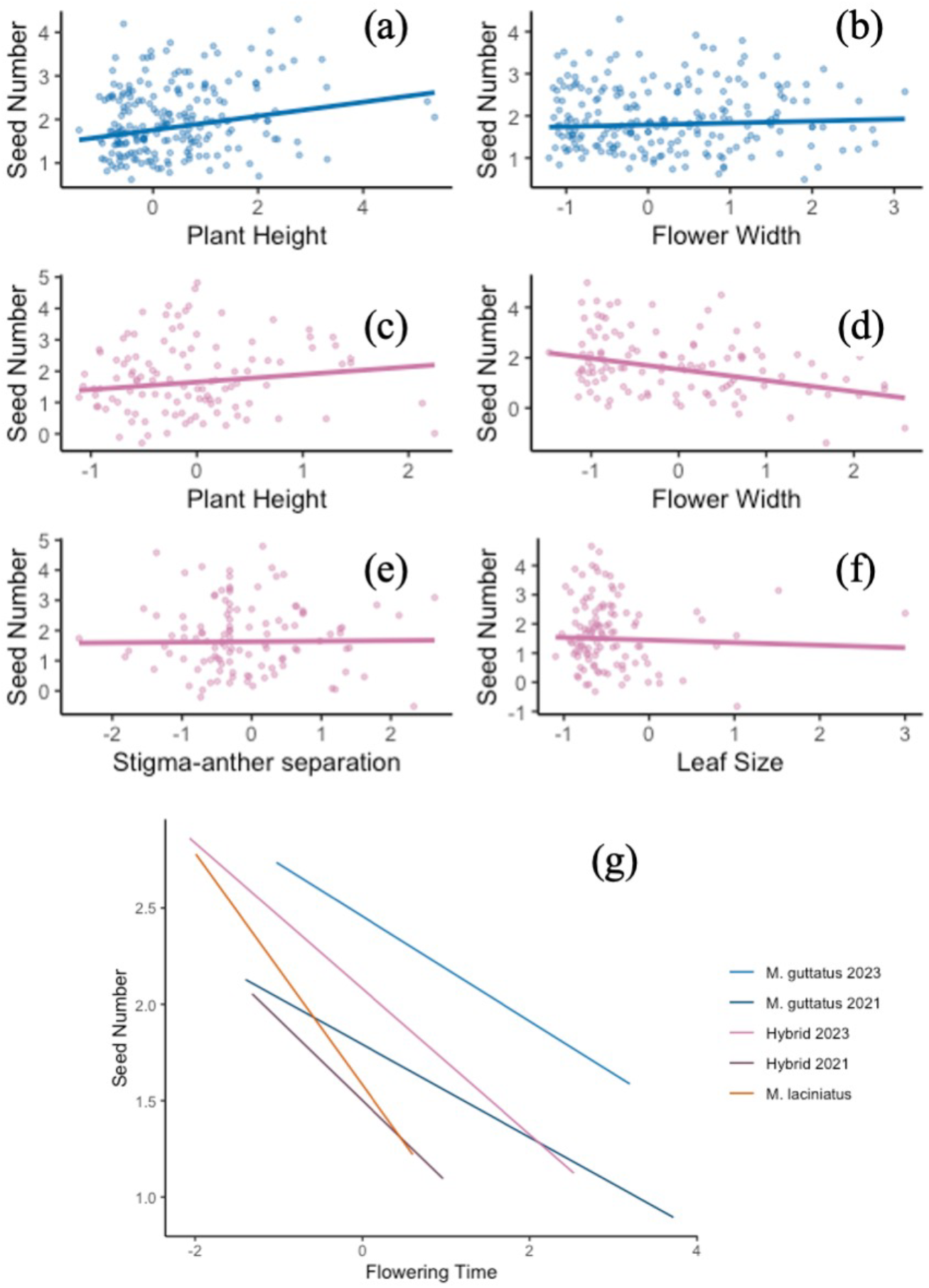
Directional selection on traits with all genotypes combined in *M. guttatus* (blue), Hybrid (pink), and *M. laciniatus* (orange) sites, visualized using partial residuals from multiple regression of best fit zero-truncated poisson models (Breheny and Burchett 2017). The fitted curves show best fitting linear regression in 2021 (a-f) and both years (g). Plant height (a, c) and flower width (b) were in the best fit models for both sites, while leaf size (c) and stigma anther separation (d) were only in best fit models for the hybrid habitat. Graph (e) also includes selection on flowering time measured in 2023 in M. guttatus (dark blue), hybrid (dark pink) and M. laciniatus (orange) habitats.

**Table 4.**
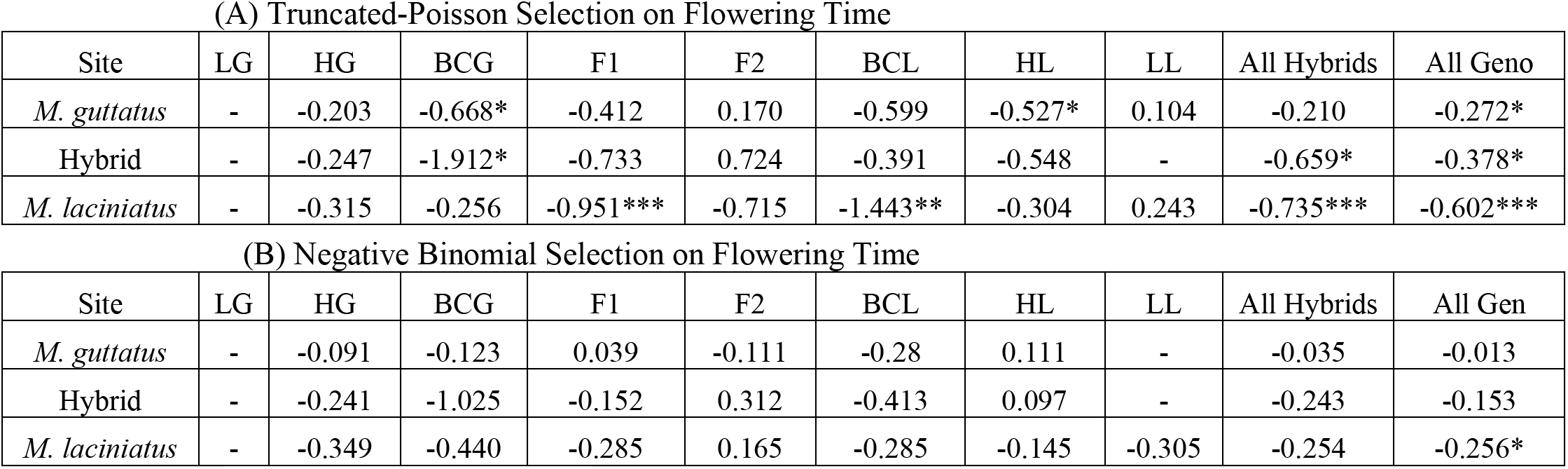
2023 selection gradients (β) on flowering time from the best fit. (A) zero-truncated poisson and (B) negative binomial models based on seed number. β indicates strength and direction of selection while asterisks indicate significance: p<.05 *, P<.01**, P< .001 ***.

(A) Truncated-Poisson Selection on Flowering Time
| Site | LG | HG | BCG | F1 | F2 | BCL | HL | LL | All Hybrids | All Geno |
| --- | --- | --- | --- | --- | --- | --- | --- | --- | --- | --- |
| *M. guttatus* | - | -0.203 | -0.668* | -0.412 | 0.170 | -0.599 | -0.527* | 0.104 | -0.210 | -0.272* |
| Hybrid | - | -0.247 | -1.912* | -0.733 | 0.724 | -0.391 | -0.548 | - | -0.659* | -0.378* |
| *M. laciniatus* | - | -0.315 | -0.256 | -0.951*** | -0.715 | -1.443** | -0.304 | 0.243 | -0.735*** | -0.602*** |

Table 4. 2023 selection gradients ( $\beta$ ) on flowering time from the best fit (A) zero-truncated poisson and (B) negative binomial models based on seed number.
| Site | LG | HG | BCG | F1 | F2 | BCL | HL | LL | All Hybrids | All Gen |
| --- | --- | --- | --- | --- | --- | --- | --- | --- | --- | --- |
| *M. guttatus* | - | -0.091 | -0.123 | 0.039 | -0.111 | -0.28 | 0.111 | - | -0.035 | -0.013 |
| Hybrid | - | -0.241 | -1.025 | -0.152 | 0.312 | -0.413 | 0.097 | - | -0.243 | -0.153 |
| *M. laciniatus* | - | -0.349 | -0.440 | -0.285 | 0.165 | -0.285 | -0.145 | -0.305 | -0.254 | -0.256* |

### Discussion

The contribution of hybridization to adaptation and speciation has long interested evolutionary biologists. To investigate the evolutionary forces maintaining hybrid zones between two sympatric Monkeyflower species, we performed repeated reciprocal transplants between native hybrid and pure species’ habitats. We predicted that hybridization is maintained in *M. guttatus-M. laciniatus* hybrid zones by a hybrid fitness advantage, while parental species’ habitats have decreased hybrid fitness and increased fitness of local genotypes (Figure 2). We instead found relaxation of selection on hybrids in the hybrid zone in both years and local maladaptation in the *M. guttatus* habitat. There was evidence of temporally varying positive and negative selection acting on hybrids, with selection for *M. laciniatus*-like traits in the hybrid zone in the dry year, and fitness advantage of F_1_ hybrids but expression of genetic incompatibilities in F_2_ hybrids in the wet year. Identifying how hybridization interacts with natural selection is essential for understanding how plant populations will persist in future changing conditions.

### A shifting hybrid zone

Hybridization might be adaptive in the wild if the hybrid zone is environmentally intermediate between parental species’ habitats. To identify whether our hybrid zone was environmentally intermediate between *M. laciniatus* and *M. guttatus’s* habitats, we compared fine-scale environmental variation and its effect on plant fitness between sites. Soil moisture decrease was strongly associated with plant survival in our experiments, confirming findings of previous studies in this system (Ferris and Willis 2018, Tataru, et al. 2023). The hybrid site had soil moisture levels and curves that fluctuated between years and were either intermediate to parental sites (dry year), or more similar to *M. laciniatus’* rocky outcrop habitat (wet year).

Temporal variation in differentiation between hybrid and parental species’ environments and equal fitness of hybrids and parental species in hybrid zones has also been found in Louisiana *Irises* (Emms and Arnold 1997). While hybrid zones as clines between parental species’ habitats have been well described (reviewed in Arnold, 1997), not many studies have investigated how temporal variation in environmental variables effect the maintenance of hybridization. Although soil moisture was sometimes more similar to *M. laciniatus’s* rocky habitat, high herbivory levels in the hybrid zone in both years were more similar to *M. guttatus’s* meadow habitat and the combination of these divergent environmental pressures could cause a hybrid’s unique blend of parental alleles to be adaptive. Examining the influence of multiple kinds of environmental variation constructs a more holistic picture of whether a hybrid zone is truly ecologically intermediate or unique compared to parental habitats.

### Relaxed selection against hybrids in hybrid zone

Given that our hybrid habitat was environmentally intermediate in some years we next sought to identify whether hybrids had a fitness advantage in the hybrid habitat. Instead of clear selection for hybridization, we found relaxation of selection against hybrids in the hybrid habitat in both years, with the exception of high F_1_ fitness in the wet year. Surprisingly, we did not find an advantage of local genotypes in each parental habitat. Instead, *M. laciniatus* had higher total fitness than *M. guttatus* in *M. guttatus’s* habitat in both years (Figure 4). In the extremely high snowpack year, non-local *M. laciniatus* had significantly higher fitness in the *M. guttatus* meadow habitat than in its own rocky outcrop environment. This asymmetry in parental fitness suggests that *M. laciniatus* is better adapted to both species habitats and could indicate a shifting fitness landscape in *M. guttatus’s* meadows due to recent climate change.

Relaxed selection against hybrids may be due to the environmentally intermediate nature of our hybrid zone (Figure 3). A mixture of drought-adapted *M. laciniatus* alleles with more mesic *M. guttatus* genetic background could have similar fitness to parental species over time in a hybrid environment with soil moisture levels that fluctuate between the two species’ habitats. In dry years like 2021 it is potentially adaptive to have a higher proportion of drought adapted-*M. laciniatus* genetic background as seen by BCL hybrids having the highest fitness and BCG having higher fitness than pure *M. guttatus* in the hybrid zone (Figure 4b). This pattern of *M. laciniatus* alleles being advantageous in an *M. guttatus* background would be consistent with patterns of asymmetric introgression in other self-fertilizing-outcrossing species pairs. Genomic analyses of gene flow between *Mimulus nasutus* and *M. guttatus* found asymmetric and recurring introgression of the drought-adapted self-fertilizing species into *M. guttatus* (Kenney and

Sweigart 2016). This same pattern was detected between *Clarkia xantiana* (self-fertilizing) and *Clarkia parviflora* (outcrossing) as well as evidence of increased introgression between species when spring precipitation is more variable (Sianta, et al. 2024).

### Fluctuating selection influences hybrid fitness

We found that the strength of selection fluctuates temporally in a *M. laciniatus*-*M. guttatus* hybrid zone. The wetter year transplant had higher overall fitness than the drier year, and fitness differences between genotypes varied temporally with F_1_ hybrids having an advantage in the wetter year. Our findings of an environmentally dependent hybrid advantage (Figure 4) parallel other findings of strong GxE interactions in a greenhouse common garden with differing watering regimes of Louisiana *Iris* species and hybrids (Johnston, et al. 2001). Also in the wet year, a larger proportion of local *M. guttatus* survived to the end of the season in the *M. guttatus* habitat but never flowered, suggesting a shift in life history expression. In populations of *Streptanthus tortuousus* with variable life history expression, plants with later germination timing are more likely to perennate (Gremer, et al. 2020). While we planted cotyledons (not seeds) due to logistical constraints, we planted our seedlings later in the high snowpack year which could have led to a proportion of the *M. guttatus*, which are known to be facultatively perennial at high elevations (Friedman and Willis 2013), behaving as perennials. Our findings demonstrate how annual environmental variation plays an important role in life history cues and subsequent population structure.

Another observed fluctuation in genotypic selection was that in the wetter year we found low F_2_ relative to F_1_ fitness in each habitat, but similar fitness between the two hybrid generations in the dry year (Figure 4). This pattern of low F_2_ relative fitness is consistent with the expression of genetic incompatibilities between the two species, that are masked in the heterozygous F_1_ but expressed in later generation hybrids (Coyne and Orr 2004). Previous studies of greenhouse crosses between *M. guttatus* and *M. laciniatus* have not found hybrid breakdown (Vickery 1964), suggesting that the expression of these BDMIs may be environmentally dependent. Environmental dependence, or extrinsic post-zygotic reproductive isolation, is consistent with the fitness difference only appearing in the wet year. A field transplant with multiple ecotypes and hybrids of *Senecio lautus* also found an environmentally dependent decrease in F_2_ relative to F_1_ fitness (Walter, et al. 2020). The environmentally dependent expression of BDMIs in later generation hybrids might balance any selective advantage experienced by early generation hybrids or drought adapted-*M. laciniatus* alleles in the hybrid zone.

### Drought adapted traits are advantageous in the hybrid zone

*M. laciniatus* is adapted to harsh and ephemeral habitats, so genetic material from this species may help with adaptation to dry conditions like those seen in 2021 and sometimes in the hybrid zone (Figure 3). The direction of phenotypic selection in the hybrid zone was largely towards *M. laciniatus*-like trait values, with the strongest selection on traits involved in reproductive isolation: flowering time and flower size. Adaptive introgression in hybrids has been demonstrated in a number of systems (Seehausen 2004) such as in *Helianthus* where there has been adaptative introgression of abiotic tolerance (Whitney, et al. 2010) and herbivory resistance (Whitney, et al. 2006). Subsequent experimental Sunflower hybrid transplants found increased hybrid fitness and faster trait evolution in hybrid than non-hybrid populations (Mitchell, et al. 2019). This suggests that patterns of selection for *M. laciniatus*-like traits in the hybrid zone are a signature of adaptive introgression.

## Conclusions

Our findings suggest that in this system hybridization is maintained by relaxed selection against hybridization in the hybrid zone and temporally varying selection (Arnold 1997). Gene flow between our incompletely reproductively isolated species may allow for advantageous introgression of drought-adapted traits from a rocky outcrop specialist, *M. laciniatus,* into the more mesic-adapted *M. guttatus* in drier years. This is possibly due to the hybrid zone’s being environmentally intermediate in some ways. Novel combinations of traits in our hybrids paired with strong GxE interactions may provide a fitness benefit, however BDMIs expressed in later generation hybrids might limit this benefit in years with high water availability, a form of migration-selection balance (Key 1968). While selection varied temporally in one hybrid zone, identifying whether selection varies across multiple *Mimulus* hybrid zones would broaden understanding of the maintenance of hybridization, and how patterns of gene flow vary across space as well as time. Overall, our study demonstrates that large-scale reciprocal transplants are important in identifying how variation in natural selection impacts the maintenance of natural hybrid zones.

## Author contributions

DT designed experiment, conducted all greenhouse and field work, data analysis, and wrote the manuscript (original draft, review, and editing). AER, FM, ML, and SD all assisted with field work and data processing. KGF assisted with experimental conceptualization, funding acquisition, and writing (review and editing).

## Acknowledgments

Thank you to the UCNRS Yosemite Field Station for housing and research support in both years of the experiment (doi:10.21973/N3V36C), and the Yosemite National Park Service for research support and permitting (Permit #YOSE-2021-SCI-0034 and YOSE-2023-SCI-0018). Thank you also to Caroline Dong, Bolivar Aponte-Rolon, Juj Sullivan, Grace Fitzgerald-Diaz, and Jessica Scales for help with plant outs. Thank you to the undergraduate students and technicians who counted seeds, helped with grow-outs, and analyzed leaf shape; Amanda Augustino, Charlotte Hankin, Kennedy Derosin, and Sarah Uher.

We acknowledge the long-standing stewards of the unceded land upon which our field work was conducted, the Southern Sierra Miwuk Nation, Bishop Paiute Tribe, Bridgeport Indian Colony, Mono Lake Kutzadikaa, North Fork Rancheria of Mono Indians of California, Picayune Rancheria of the Chukchansi Indians and the Tuolumne Band of Me-Wuk Indians. As scientists, we strive to take responsibility for the impacts of colonialism in our field and move forward with respect and support of indigenous movements and knowledge.

## Data and code availability statement

Data and code available from the Dryad Digital Repository.

## Works Cited

1. Abbott, R, D Albach, S Ansell, J Arntzen, S Baird, N Bierne, J Boughman, et al. 2013. “Hybridization and speciation.” Journal of Evolutionary Biology 26 (2): 229–246.

2. Anderson, E, and G L Stebbins. 1954. “Hybridization as an Evolutionary Stimulus.” Evolution 8 (4): 378–388.

3. Arnold, ML. 1997. Natural hybridization and evolution. New York: Oxford University Press.

4. Bartoń, Kamil. 2009. Mu-MIn: Multi-model inference. http://R-Forge.R-project.org/projects/mumin/.

5. Barton, NH, and G M Hewitt. 1985. “Analysis of Hybrid Zones.” Annual Review of Ecology and Systematics 16: 113–148.

6. Bateson, W. 1909. “Heredity and variation in modern lights.” In Darwin and Modern Science, by AC Seward, 85–101. Cambridge UK: Campbridge University Press.

7. Brandvain, Y, A Kenney, L Flagel, G Coop, and A Sweigart. 2014. “Speciation and introgression between Mimulus nasutus and Mimulus guttatus.” PLoS Genetics 10 (6): e1004410.

8. Brasier, CM. 1995. “Episodic selection as a force in fungal microevolution, with special reference to clonal speciation and hybrid introgression.” Canadian Journal of Botany 73: S1213–S1221.

9. Breheny, Patrick, and Woodrow Burchett. 2017. “Visualization of Regression Models Using visreg.” The R Journal 9 (2): 56–71.

10. Brooks, ME, K J van Benthem, A Magnusson, C W Berg, H J Skaug, M Maechler, and B M Bolker. 2017. “glmmTMB Balances Speed and Flexibility Among Packages for Zero-inflated Generalized Linear Mixed Modeling.” The R Journal 9 (2): 378–400.

11. Campbell, D, J Powers, and M Crowell. 2024. “Pollinator and habitat-mediated selection as potential contributors to ecological speciation in two closely related species.” Evolution Letters 8 (2): 311–321.

12. Canty, A, and B D Rippley. 2024. “boot: Bootstrap R (S-Plus) Functions.” R package version 1.3-30.

13. CDEC. 2024. Data for average snowpack. May 6. https://cdec.water.ca.gov/snowapp/sweq.action.

14. Coyne, JA, and H A Orr. 2004. Speciation. Sunderland, MA: Sinauer Associates.

15. Cuthill, IC, AT D Bennett, J C Partridge, and E J Maier. 1999. “Plumage Reflectance and the Objective Assessment of Avian Sexual Dichromatism.” The American Naturalist 153 (1): 183–200.

16. Davison, AC, and D V Hinkey. 1997. Bootstrap Methods and Their Applications. Cambridge: Cambridge University Press.

17. Dobzhansky, T. 1936. “Studies on hybrid sterility. ii. Localization of sterility factors in Drosophila pseudoobscura hybrids.” Genetics 21: 113–135.

18. Emms, SK, and M L Arnold. 1997. “The effect of habitat on parental and hybrid fitness: transplant experiments with Louisiana Irises.” Evolutiopn 51 (4): 1112–1119.

19. Ferris, KG, and J H Willis. 2018. “Differential adaptation to a harsh granite outcrop habitat between sympatric Mimulus species.” Evolution 72: 1225–1241.

20. Ferris, KG, and J H Willis. 2018. “Differential adaptation to a harsh granite outcrop habitat between sympatric Mimulus species.” Evolution 72: 1225–1241.

21. Ferris, KG, J P Sexton, and J H Willis. 2014. “Speciation on a local geographic scale: the evolution of a rare rock outcrop specialist in Mimulus.” The Philosophical Transactions of the Royal Society B 369: 20140001.

22. Ferris, KG, L L Barnett, B K Blackman, and J H Willis. 2017. “The genetic architecture of local adaptation and reproductive isolation in sympatry within the Mimulus guttatus species complex.” Molecular Ecology 26 (1): 208–224.

23. Ferris, KG, T Rushton, A B Grenenlee, K Toll, B K Blackman, and J H Willis. 2015. “Leaf shape evolution has a similar genetic architecture in three edaphic specialists within the Mimulus guttatus species complex.” Annals of Botany 116: 213–223.

24. Friedman, J, and J H Willis. 2013. “Major QTLs for critical photoperiod and vernalization underlie extensive variation in flowering in the Mimulus guttatus species complex.” New Phytologist 199: 571–583.

25. Gay, L, P Crotchet, D Bell, and T Lenmormand. 2008. “Comparing clines on molecular and phenotypic traits in hybrid zones: A window on tension zone models.” Evolution 62 (11): 2789–2806.

26. Grant, BR, and P R Grant. 1996. “Cultural inheritance of song and its role in evolution of Darwin’s finches.” Evolution 50 (6): 2471–2478.

27. Grant, PR, and B R Grant. 2016. “Introgressive hybridization and natural selection in Darwin’s finches.” Biological Journal of the Linnean Society 117 (4): 812–822.

28. Grant, V. 1966. “The selective origin of incompatability barriers in the plant genus gilia.” The American Naturalist 100 (911): 99–118.

29. Gremer, J, c Wilcox, A Chiono, E Suglia, and J Schmitt. 2020. “Germination timing and chilling exposure create contingency in life history and influence fitness in the native wildflower Streptanthus tortuosus.” Journal of Ecology 108 (1): 239–255.

30. Hall, MC, and J H Willis. 2006. “Divergent selection on flowering time contributes to local adaptation in Mimulus guttatus populations.” Evolution 60: 2466–2477.

31. Hatfield, T, and D Schluter. 1999. “Ecological speciation in sticklebacks: environment-dependent hybrid fitness.” Evolution 53 (3): 866–873.

32. Johnston, JA, D J Grise, L A Donovan, and M L Arnold. 2001. “Environment-dependent performance and fitness of Iris brevicaulis, I. fulva (Iridaceae), and hybrids.” American Journal of Botany 88 (5): 933–938.

33. Kenney, A, and A Sweigart. 2016. “Reproductive isolation and introgression between sympatric Mimulus species.” Molecular ecology 25 (11): 2499–2517.

34. Key, KHL. 1968. “The concept of stasipatric speciation.” Systematic Zoology 14–22.

35. Kim, S, and L H Rieseberg. 1999. “Genetic Architecture of Species Differences in Annual Sunflowers: Implications for Adaptive Trait Introgression.” Genetics 153: 965–977.

36. Kingsolver, JG, H E Hoekstra, J M Hoekstra, D Berrigan, S N Vignieri, C E Hill, A Hoang, P Gilbert, and P Beerli. 2001. “The strength of phenotypic selection in natural populations.” The American Naturalist 157 (3): 245–261.

37. Lande, R, and S J Arnold. 1983. “The measurement of selection on correlated characters.” Evolution 37: 1210–1226.

38. Lenth, RV. 2016. “Least-Squares Means: The R Package lsmeans.” Journal of Statistical Software 69 (1): 1–33.

39. Lexer, C, L H Rieseberg, and R A Randell. 2003. “Experimental hybridization as a tool for studying selection in the wild.” Ecology 84 (7): 1688–1699.

40. Mayr, E. 1942. Systematics and the Origin of Species. New York: Columbia University Press.

41. Miglia, KJ, E D McArthur, W S Moore, H Wang, J H Graham, and D C Freeman. 2005. “Nine-year reciprocal transplant experiment in the gardens of the basin and mountain big sagebrush (Artemisia tridentata: Asteraceae) hybrid zone of Salt Creek Canyon: the importance of multiple-year tracking of fitness.” Biological Journal of the Linnean Society 86: 213–225.

42. Mitchell, N, G Owens, S Hovick, L H Rieseberg, and K Whitney. 2019. “Hybridization speeds adaptive evolution in an eight-year field experiment.” Scientific Reports 9 (1): 1–12.

43. Mitchell-Olds, T, and R G Shaw. 1987. “Regression analysis of natural selection: statistical inference and biological interpretation.” Evolution 41: 1149–1161.

44. Moore, WS. 1977. “An evaluation of narrow hybrid zones in vertebrates.” The Quarterly Review of Biology 263–277.

45. Moran, BM, C Payne, Q Langdon, D L Powell, Y Brandvain, and M Schumer. 2021. “The genomic consequences of hybridization.” eLife 10: e69016.

46. Muller, HJ. 1942. “Isolating mechanisms, evolution and temperature.” Biology Symposium 71–125.

47. Pinheiro, J, D Bates, and R Core Team. 2023. nlme: Linear and Nonlinear Mixed Effects Models. https://CRAN.R-project.org/package=nlme.

48. Ruhsam, M, P M Hollingsworth, and R A Ennos. 2011. “Early evolution in a hybrid swarm between outcrossing and selfing lineages in Geum.” Heredity 107: 246–255.

49. Schemske, DW. 2000. “Review: Understanding the Origin of Species.” Evolution 54 (3): 1069-1073.

50. Seehausen, O. 2004. “Hybridization and adaptive radiation.” Trends in Ecology and Evolution 19 (4): 198–207.

51. Sianta, SA, D A Moeller, and Y Brandvain. 2024. “The extent of introgression between incipient Clarkia species is determined by temporal environmental variation and mating system.” PNAS 121 (12): e2316008121.

52. Siepielski, AM, K M Gotanda, M B Morrissey, S E Diamond, J D Dibattista, and S M Carlson. 2013. “The spatial patterns of directional phenotypic selection.” Ecology Letters 16: 1382–1392.

53. Smith, K, H J Kearney, J Austin, and J Melville. 2013. “Molecular patterns of introgression in a classic hybrid zone between the Australian tree frogs, Litoria ewingii and L. paraewingi: Evidence of a tension zone.” Molecular Ecology 22 (7): 1869–1883.

54. Stebbins, GL. 1959. “The role of hybridizaiton in evolution.” Proceedings of the American Philosophical Society 103 (2): 231–251.

55. Svensson, Erik I. 2023. “Phenotypic selection in natural populations: what have we learned in 40 years?” Evolution 77 (7): 1492–1504.

56. Tataru, D, E C Wheeler, and K G Ferris. 2023. “Spatially and temporally varying selection influence species boundaries in two sympatric Mimulus.” Proceedings of the Royal Society B 290: 20222279.

57. Team, R Core. 2022. R: A language and environment for statistical computing. v4.2.1. Vienna. Austria: R Foundation for Statistical Computing.

58. Vickery, RK. 1964. “Barriers to gene exchange between members of the Mimulus guttatus complex (Scrophulariaceae).” Evolution 18: 52–69.

59. Wadgymar, S, S Daws, and J Anderson. 2017. “Integrating viability and fecundity selection to illuminate the adaptive nature of genetic clines.” Evolution Letters 1 (1): 26–39.

60. Walter, GM, T J Richards, M J Wilkinson, M W Blows, J D Aguirre, and D Ortiz-Barrientos. 2020. “Loss of ecologically important genetic variation in late generation hybrids reveals links between adaptation and speciation.” Evolution Letters 4 (4): 302–316.

61. Wang, H, E D McArthur, S C Sanderson, J H Graham, and D C Freeman. 1997. “Narrow hybrid zone between two subspecies of big sagebrush (Artemisia tridentata: Asteraceae). IV. Reciprocal transplant experiments.” Evolution 51 (1): 95–102.

62. Wei, Taiyun, and Viliam Simko. 2021. R package ‘corrplot’: Visualization of a Correlation Matrix (Version 0.92). https://github.com/taiyun/corrplot.

63. Whitney, KD, R A Randell, and L H Riesberg. 2010. “Adaptive introgression of abiotic tolerance traits in the sunflower Helianthus annus.” New Phytologist 187: 230–239.

64. Whitney, KG, R A Randell, and L H Rieseberg. 2006. “Adaptive introgression of herbivore resistance traits in the weedy sunflower Helianthus annus.” The American Naturalist 167: 794–807.

