## Supplemental Figures and Tables for "Fluctuating selection in a Monkeyflower hybrid zone"

***Supplementary Figures***

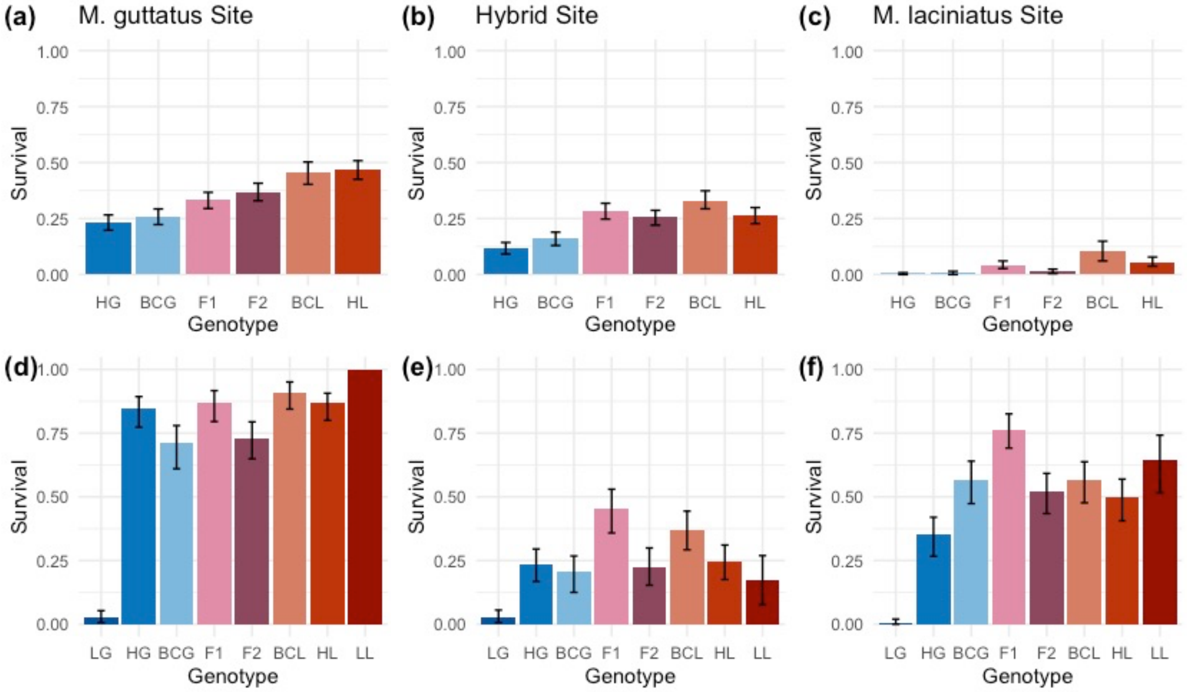


S1. Average proportion of plants that survived to flowering per genotype in each site, with sites plotted separately. Upper row (a, b, c) is 2021 sites and lower row (d, e, f) is 2023 sites. Genotypes are, from left to right, local *M. guttatus* (LG), hybrid habitat *M. guttatus* (HG), back-crossed *M. guttatus* (BCG), first generation hybrid ($F_1$), second generation hybrid ($F_2$), backcrossed *M. laciniatus* (BCL), hybrid habitat *M. laciniatus* (HL) and local *M. laciniatus* (LL). Error bars are calculated using 95% confidence intervals from bootstrap analysis.

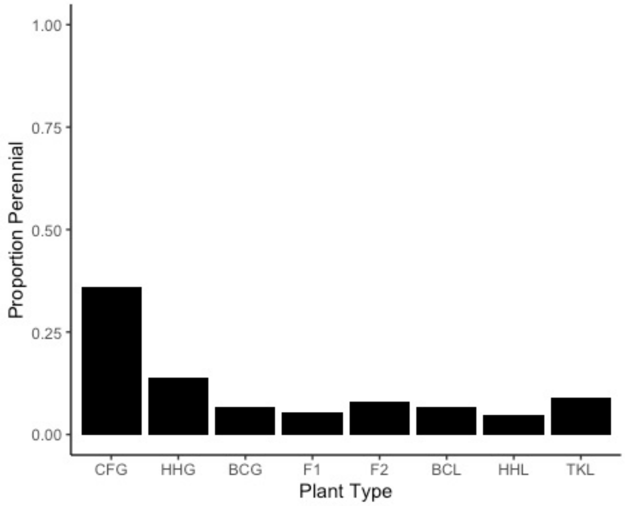


S2. Proportion of individuals planted in the *M. guttatus* site in 2023 that survived until the end of the season without flowering (proxy for perenniality).

| | Flowering Tin | Leaf Lobing | Flower Width | Plant Height | Stigma Anther Sep. | Leaf Area |
|---|---|---|---|---|---|---|
| Flowering Time | **1.00** | | | | | |
| Leaf Lobing | **0.16** | **1.00** | | | | |
| Flower Width | **0.15** | **-0.10** | **1.00** | | | |
| Plant Height | **0.24** | **-0.07** | **0.58** | **1.00** | | |
| Stigma Anther Sep. | **0.15** | **-0.04** | **0.36** | **0.26** | **1.00** | |
| Leaf Area | **-0.20** | **-0.15** | **0.42** | **0.41** | **0.20** | **1.00** |

S3. Trait correlations of standardized traits for all genotypes and sites in 2021. Correlations did not vary significantly between sites.

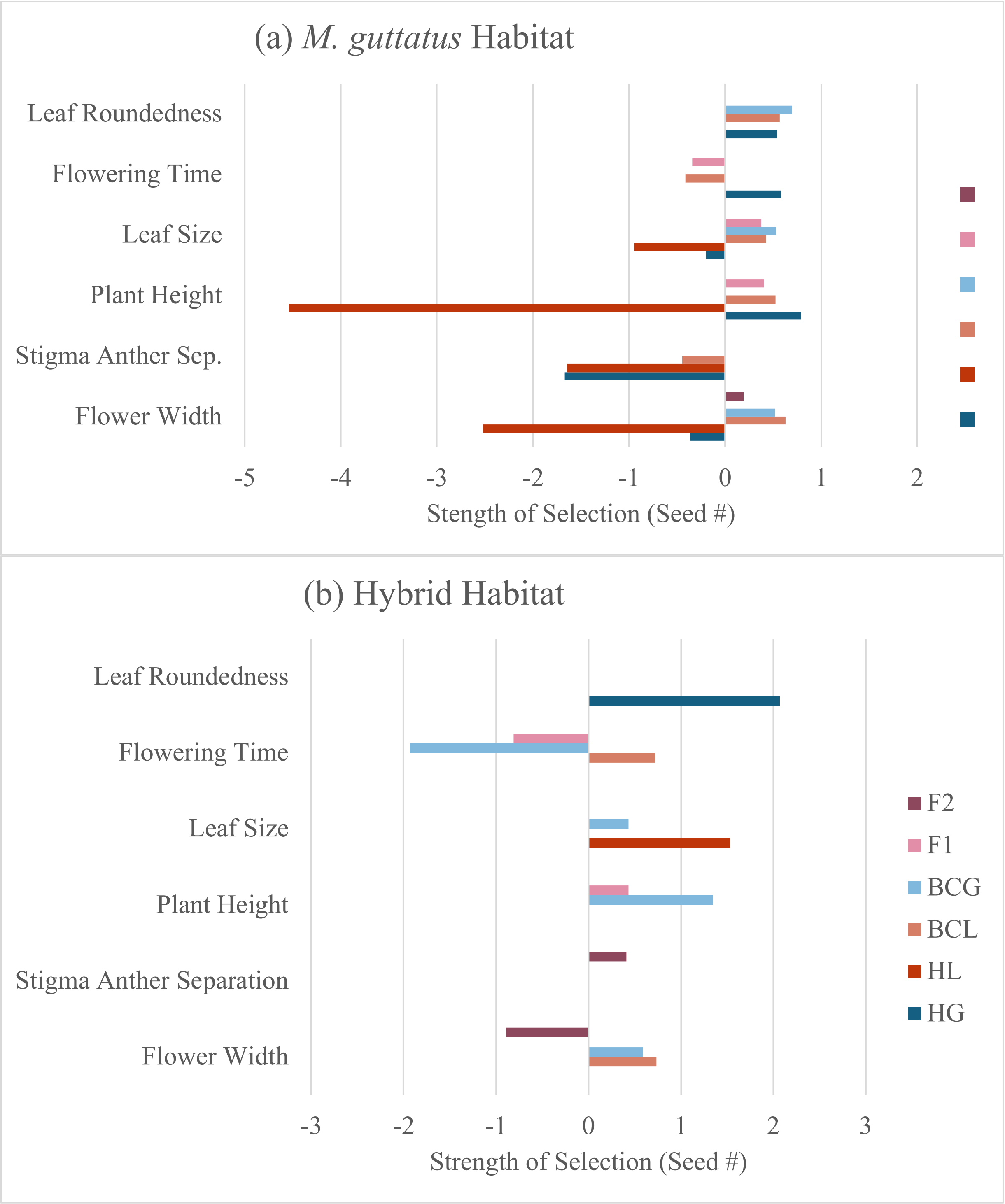


S4. Phenotypic selection gradients from zero-truncated poisson models of the 2021 transplant in the *M. guttatus* (a) and hybrid (b) sites. Values from these graphs can be found in Table 3.

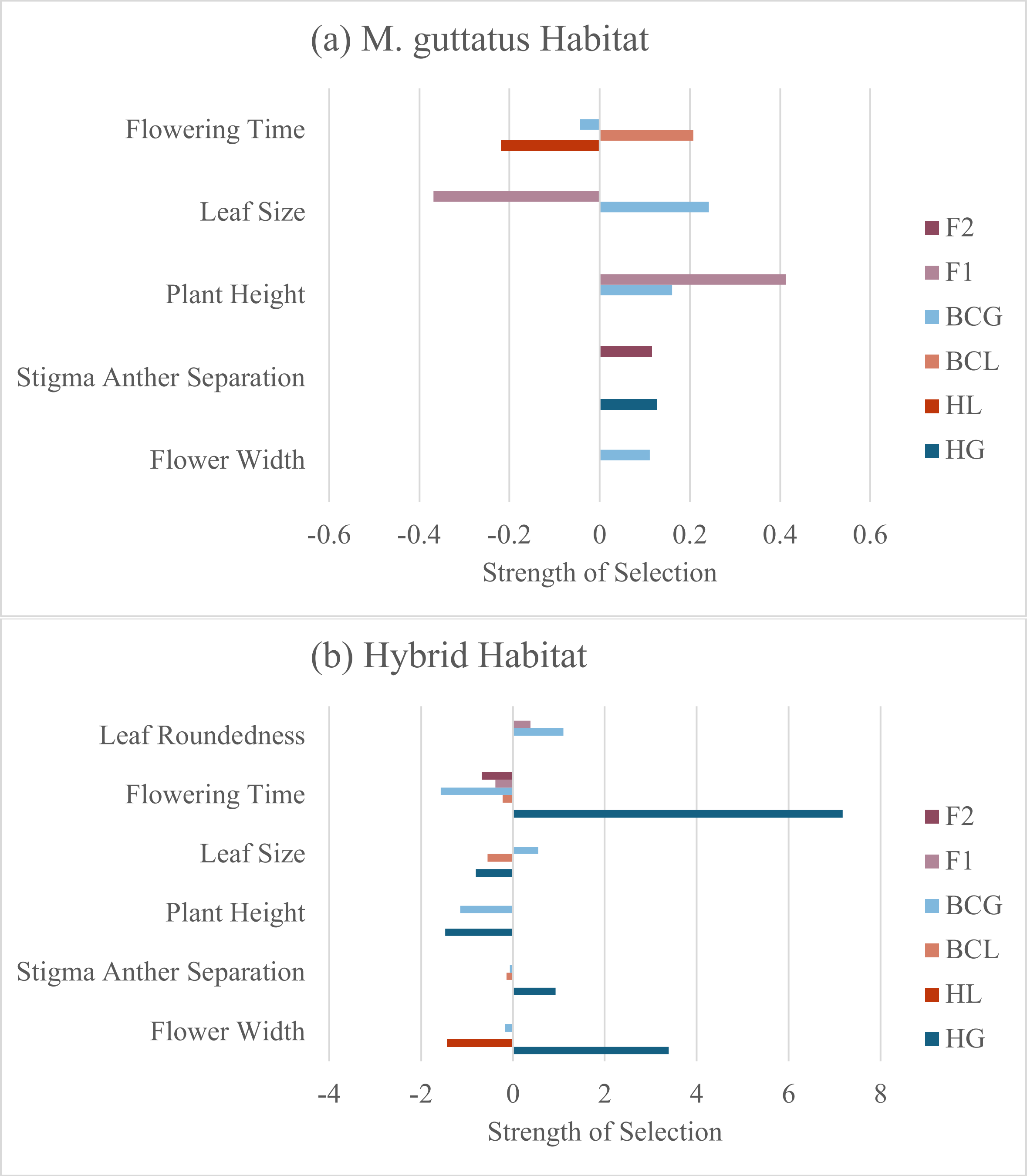


S5. Phenotypic selection gradients from negative binomial of the 2021 transplant in the *M. guttatus* (a) and hybrid (b) sites. Values from these graphs can be found in supplementary table S4.

***Supplementary Tables***

| Year | Site | Genotype | # Planted | Date Planted | Avg. Fecundity | 95% CI |
|---|---|---|---|---|---|---|
| 2021 | *M. guttatus* | HG | 600 | 4/28/21 | 0.537 | 0.3527, 0.81 |
| 2021 | *M. guttatus* | BCG | 600 | 4/28/21 | 0.604 | 0.369, 0.925 |
| 2021 | *M. guttatus* | F1 | 600 | 4/28/21 | 0.573 | 0.364, 0.864 |
| 2021 | *M. guttatus* | F2 | 600 | 4/28/21 | 0.8 | 0.554, 1.13 |
| 2021 | *M. guttatus* | BCL | 372 | 4/28/21 | 1.826 | 1.333, 2.657 |
| 2021 | *M. guttatus* | HL | 600 | 4/28/21 | 1.685 | 1.299, 2.221 |
| 2021 | Hybrid | HG | 600 | 4/9/21 | 0.155 | 0.06, 0.431 |
| 2021 | Hybrid | BCG | 600 | 4/9/21 | 0.376 | 0.221, 0.592 |
| 2021 | Hybrid | F1 | 600 | 4/9/21 | 0.636 | 0.453, 0.925 |
| 2021 | Hybrid | F2 | 600 | 4/9/21 | 0.578 | 0.379, 0.909 |
| 2021 | Hybrid | BCL | 498 | 4/9/21 | 0.847 | 0.571, 1.181 |
| 2021 | Hybrid | HL | 600 | 4/9/21 | 0.718 | 0.527, 0.972 |
| 2021 | *M. laciniatus* | HG | 300 | 5/26/21 | 0 | NA |
| 2021 | *M. laciniatus* | BCG | 342 | 5/26/21 | 0.006 | 0, 0.018 |
| 2021 | *M. laciniatus* | F1 | 600 | 5/26/21 | 0.013 | 0, 0.05 |
| 2021 | *M. laciniatus* | F2 | 462 | 5/26/21 | 0 | NA |
| 2021 | *M. laciniatus* | BCL | 144 | 5/26/21 | 0.027 | 0, 0.069 |
| 2021 | *M. laciniatus* | HL | 510 | 5/26/21 | 0.016 | 0.002, 0.049 |
| 2023 | *M. guttatus* | LG | 150 | 6/7/23 | 0.127 | 0, 0.588 |
| 2023 | *M. guttatus* | HG | 150 | 6/7/23 | 4.233 | 2.909, 6.318 |
| 2023 | *M. guttatus* | BCG | 118 | 6/7/23 | 2.9 | 1.828, 4.489 |
| 2023 | *M. guttatus* | F1 | 138 | 6/7/23 | 7.568 | 5.249, 11.828 |
| 2023 | *M. guttatus* | F2 | 150 | 6/7/23 | 1.291 | 0.681, 2.326 |
| 2023 | *M. guttatus* | BCL | 122 | 6/7/23 | 4.549 | 3.160, 6.608 |
| 2023 | *M. guttatus* | HL | 150 | 6/7/23 | 10.567 | 8.10, 14.43 |
| 2023 | *M. guttatus* | LL | 12 | 6/7/23 | 3.909 | 0.455, 11 |
| 2023 | Hybrid | LG | 150 | 5/7/23 | 0.208 | 0.028, 0.958 |
| 2023 | Hybrid | HG | 150 | 5/7/23 | 1.342 | 0.570, 2.636 |
| 2023 | Hybrid | BCG | 150 | 5/7/23 | 1.607 | 0.727, 4.054 |
| 2023 | Hybrid | F1 | 150 | 5/7/23 | 6.157 | 3.363,11.815 |
| 2023 | Hybrid | F2 | 150 | 5/7/23 | 0.664 | 0.190, 2.423 |
| 2023 | Hybrid | BCL | 150 | 5/7/23 | 2.208 | 0.974, 6.335 |
| 2023 | Hybrid | HL | 150 | 5/7/23 | 1.838 | 1.007, 3.036 |
| 2023 | Hybrid | LL | 52 | 5/7/23 | 0.173 | 0, 0.78 |
| 2023 | *M. laciniatus* | LG | 150 | 7/13/23 | 0.188 | 0, 0.5638 |
| 2023 | *M. laciniatus* | HG | 150 | 7/13/23 | 2.04 | 1.040, 4.072 |
| 2023 | *M. laciniatus* | BCG | 150 | 7/13/23 | 2.233 | 1.353, 3.952 |
| 2023 | *M. laciniatus* | F1 | 150 | 7/13/23 | 10.705 | 8.21, 13.97 |
| 2023 | *M. laciniatus* | F2 | 150 | 7/13/23 | 3.671 | 2.404, 6.105 |
| 2023 | *M. laciniatus* | BCL | 150 | 7/13/23 | 2.819 | 1.819, 4.444 |
| 2023 | *M. laciniatus* | HL | 150 | 7/13/23 | 3.08 | 1.821, 5.212 |
| 2023 | *M. laciniatus* | LL | 62 | 7/13/23 | 5.242 | 3.484, 7.113 |

Table S1. Number of plants per genotype planted at each site in each year with date planted. Average fecundity (total seed number/number of plants planted) per genotype in each site with 95% confidence intervals from bootstrap estimates with 1000 repetitions with replacement.

| PC | Year | Site | | Block | | Pairwise Site Difference | | |
|---|---|---|---|---|---|---|---|---|
| | | *F* | *p-value* | *F* | *p-value* | H-G | L-G | L-H |
| 1 | 2021 | 211 | <2e-16 | 3.52 | 0.06 | -2.33 | -5.37 | -3.04 |
| 1 | 2023 | 555.88 | <2e-16 | 3.95 | 0.05 | -6.71 | -7.85 | -1.14 |
| 2 | 2021 | 137.9 | <2e-16 | 1.21 | 0.273 | 3.21 | 0.92 | -2.29 |
| 2 | 2023 | 120.8 | <2e-16 | 107.6 | <2e-16 | -2.52 | 0.82 | 3.34 |
| 3 | 2021 | 22.08 | 4.19e-09 | 2.94 | 0.09 | 0.43 | -0.47 | -1.22 |
| 3 | 2023 | 9.13 | 0.0002 | 1.16 | 0.28 | NS | -0.57 | -0.54 |

Table S2. Results of ANOVAs of soil moisture PCA analysis, with PC axis as dependent variable and site and block as independent variables. Pairwise site difference refers to significant differences in Tukey post hoc tests of ANOVAs.

| Independent Variable | *df* | *F statistic* | *p value* |
|---|---|---|---|
| Site | 2 | 336.262 | < 2e-16 |
| Genotype | 7 | 6.143 | 3.43e-07 |
| Year | 1 | 98.206 | < 2e-16 |
| Site x Year | 2 | 25.152 | 1.25e-11 |
| Site x Genotype | 14 | 3.426 | 1.37e-05 |
| Genotype x Year | 5 | 7.819 | 2.33e-07 |
| Site x Genotype x Year | 10 | 4.508 | 2.17e-06 |

Table S3. Results of best fit linear mixed effects models for effect of site, genotype, year, and their interaction (independent variables) on herbivory (dependent variable).

| Trait | Genotype (G) | | Environment (E) | | G x E | |
|---|---|---|---|---|---|---|
| | *F statistic* | *p value* | *F statistic* | *p value* | *F statistic* | *p value* |
| Flowering Time | 130.3 | < 0.001 | 102.1 | < 0.001 | 8.6 | < 0.001 |
| Plant Height | 110.7 | < 0.001 | 29.3 | < 0.001 | 5.2 | < 0.001 |
| Stigma-Anther Sep. | 60.4 | < 0.001 | 11.3 | < 0.001 | 2.6 | 0.007 |
| Leaf Size | 21.6 | < 0.001 | 68.7 | < 0.001 | 2.7 | 0.004 |
| Flower Size | 140.3 | < 0.001 | 18.6 | < 0.001 | 1.4 | 0.1938 |
| Leaf Lobing | 4.1 | 0.0015 | 65.5 | < 0.001 | 0.9 | 0.5237 |
| Flowering Time (2023) | 22.1 | < 0.001 | 618.6 | < 0.001 | 6.2 | < 0.001 |

Table S4. Results of linear mixed effects models of effect of genotype, environment, and their interaction (independent variables) on trait values (dependent variable) for all sites combined in 2021, and for flowering time in 2023.

(a) Hybrid Habitat

| Trait | HG | HL | BCL | BCG | F1 | F2 | All Hybrids | All Geno |
|---|---|---|---|---|---|---|---|---|
| Flower Width (FW) | 3.396 | -1.439 | - | -0.179 | - | - | - | - |
| Stigma Anther Separation (SA) | 0.927 | - | -0.144 | -0.066 | - | | - | - |
| Plant Height (PH) | -1.48 | - | - | -1.144 | - | - | - | - |
| Leaf Area (LA) | -0.805 | - | -0.549 | 0.550 | - | - | - | - |
| Flowering Time (FT) | 7.171 | - | -0.223 | -1.568 | -0.379 | -0.676 | -0.439 | -0.427 |
| Leaf Roundedness (LR) | - | - | - | 1.090 | 0.379 | - | - | - |
| FT x FW | -6.525 | - | - | 0.160 | - | - | - | - |
| FT x SA | - | - | -0.055 | - | - | - | - | - |
| FT x PH | 1.476 | - | - | -2.650 | - | - | - | - |
| FW x PH | - | - | - | 0.610 | - | - | - | - |

(b) *M. guttatus* habitat

| Trait | HG | HL | BCL | BCG | F1 | F2 | All Hybrids | All Geno |
|---|---|---|---|---|---|---|---|---|
| Flower Width (FW) | - | - | - | 0.111 | - | - | - | - |
| Stigma Anther Separation (SA) | 0.127 | - | - | - | - | 0.116 | - | - |
| Plant Height (PH) | - | - | - | 0.161 | 0.413 | - | 0.109 | - |
| Leaf Area (LA) | - | - | - | 0.242 | -0.37 | - | - | - |
| Flowering Time (FT) | - | -0.21 | 0.208 | -0.044 | - | - | - | - |
| Leaf Roundedness (LR) | - | - | - | - | - | - | -0.173 | 0.101 |
| FT x PH | - | - | - | -0.052 | - | - | - | - |
| FT x FW | - | - | - | -0.022 | - | - | - | - |

Table S5. Selection gradients (β) from the best fit negative binomial; models based on 2021 seed number in the hybrid habitat (A) and *M. guttatus* habitat (B). β indicates strength and direction of selection.
